# Mechanism of SARS-CoV-2 polymerase inhibition by remdesivir

**DOI:** 10.1101/2020.10.28.358481

**Authors:** Goran Kokic, Hauke S. Hillen, Dimitry Tegunov, Christian Dienemann, Florian Seitz, Jana Schmitzova, Lucas Farnung, Aaron Siewert, Claudia Höbartner, Patrick Cramer

## Abstract

Remdesivir is the only FDA-approved drug for the treatment of COVID-19 patients^1–4^. The active form of remdesivir acts as a nucleoside analogue and inhibits the RNA-dependent RNA polymerase (RdRp) of coronaviruses including SARS-CoV-2^5–7^. Remdesivir is incorporated by the RdRp into the growing RNA product and allows for addition of three more nucleotides before RNA synthesis stalls^6,8^. Here we use synthetic RNA chemistry, biochemistry and cryo-electron microscopy to establish the molecular mechanism of remdesivir-induced RdRp stalling. We show that addition of the fourth nucleotide following remdesivir incorporation into the RNA product is impaired by a barrier to further RNA translocation. This translocation barrier causes retention of the RNA 3’-nucleotide in the substrate-binding site of the RdRp and interferes with entry of the next nucleoside triphosphate, thereby stalling RdRp. In the structure of the remdesivir-stalled state, the 3’-nucleotide of the RNA product is matched with the template base, and this may prevent proofreading by the viral 3’-exonuclease that recognizes mismatches^9,10^. These mechanistic insights should facilitate the quest for improved antivirals that target coronavirus replication.

---

Coronaviruses use an RdRp enzyme to carry out replication and transcription of their RNA genome^11–15^. The RdRp consists of three non-structural protein (nsp) subunits, the catalytic subunit nsp1216 and the accessory subunits nsp8 and nsp7^13,17^. Structures of the RdRp of SARS-CoV-2 were obtained in free form18 and with RNA template-product duplexes^8,19,20^. Together with a prior structure of SARS-CoV RdRp^21^, these results have elucidated the RdRp mechanism. For RNA-dependent RNA elongation, the 3’-terminal nucleotide of the RNA product chain resides in the –1 site and the incoming nucleoside triphosphate (NTP) substrate binds to the adjacent +1 site. Catalytic nucleotide incorporation then triggers RNA translocation and liberates the +1 site for binding of the next incoming NTP.

The nucleoside analogue remdesivir inhibits the RdRp of coronaviruses^5–7,22,23^ and shows antiviral activity in cell culture and animals^17,24–26^. Remdesivir is a phosphoramidate prodrug that is metabolized in cells to yield an active NTP analogue7 that we refer to as remdesivir triphosphate (RTP). Biochemical studies showed that the RdRp can use RTP as a substrate, leading to the incorporation of remdesivir monophosphate (RMP) into the growing RNA product^6,8^. After RMP incorporation, the RdRp extends RNA by three more nucleotides before it stalls^6,8^. This stalling mechanism appears to be specific to coronaviruses because the RdRp of Ebola virus can add five RNA nucleotides after RMP incorporation before it stalls^27^.

Recent structural studies trapped RdRp-RNA complexes after remdesivir addition to the RNA product 3’-end. One structure contained RMP in the +1 site^19^, whereas another structure contained RMP in the –1 site^8^. In both structures, RMP mimics adenosine monophosphate (AMP) and forms standard Wat-son-Crick base pairs with uridine monophosphate (UMP) in the RNA template strand. Thus, these studies explained how RMP is incorporated into RNA instead of AMP. However, they do not explain how remdesivir inhibits the RdRp because RdRp stalling occurs only after three more nucleotides have been added to the RNA^6,8^.

To investigate remdesivir-induced RdRp stalling, we first investigated how RTP (**Figure 1a**) influences RdRp elongation activity on an RNA template-product scaffold (**Figure 1b**) using a highly defined biochemical system (**Methods**). Consistent with a recent study^6^, we observed that RMP is readily incorporated into RNA and that the RNA is subsequently elongated by three more nucleotides before the RdRp stalls (**Figure 1c, d**). At high NTP concentrations, RdRp stalling was overcome and the fulllength RNA product was formed despite the presence of RMP in the RNA product (**Figure 1c, d**). Thus remdesivir neither acts as a chain terminator nor as an elongation block, but rather triggers delayed RdRp stalling by an unknown mechanism.

**Figure 1.**
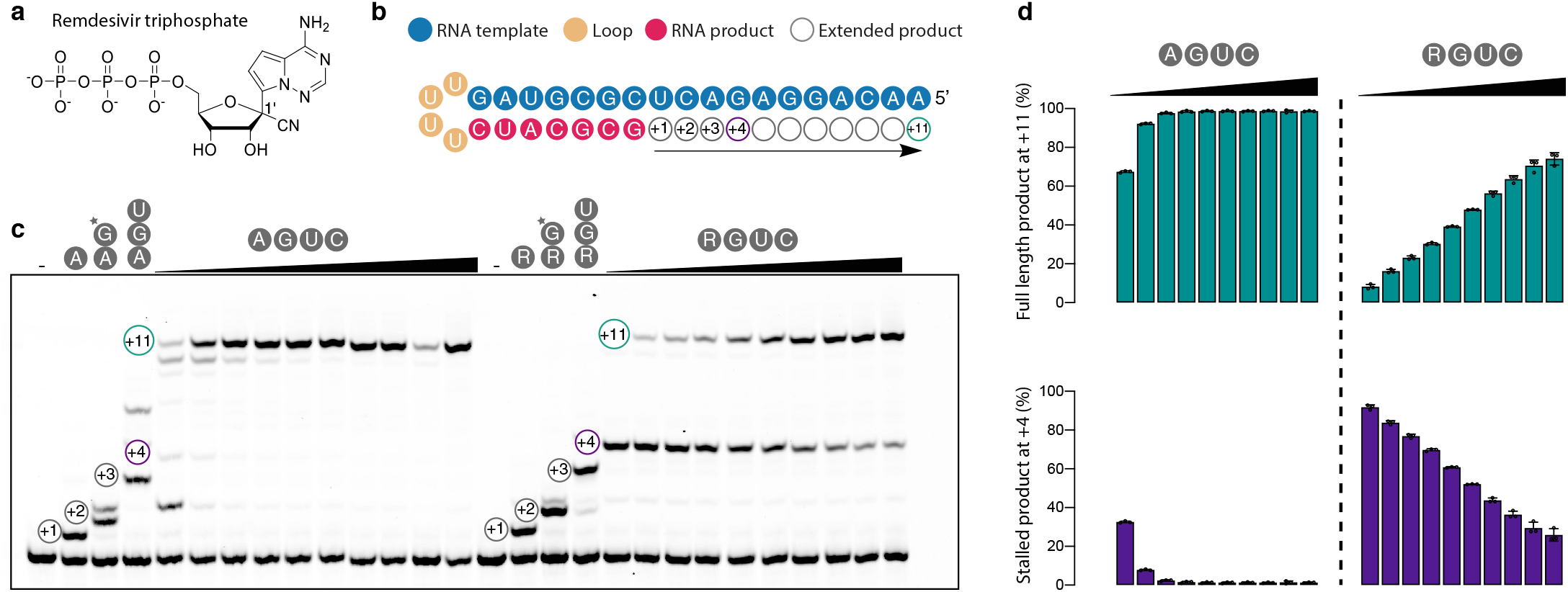
Remdesivir impairs RNA elongation by RdRp. **a** Chemical structure of remdesivir triphosphate (RTP) showing the ribose 1’ cyano group. **b** RNA template-product duplex. The direction of RNA elongation is indicated. **c** Remdesivir-induced RdRp stalling. Replacing ATP with RTP leads to an elongation barrier after addition of three more nucleotides. The barrier can be overcome at higher NTP concentrations. The RNA 5’-end contains a fluorescent label. * indicates 3’-dGTP. **d** Quantification of the experiment in panel c after triplicate measurements. Standard deviations are shown.

To uncover the mechanistic basis of remdesivir-induced RdRp stalling, we aimed to determine structures of RdRp-RNA complexes containing RMP at defined positions in the RNA product strand. We prepared RMP-containing RNA oligonucleotides by solid-phase synthesis using 5’-*O*-DMT-2’-*O*-TBDMS-protected 3’-cyanoethyl diisopropyl phosphoramidite (Rem-PA), which we synthesized from 1’-cyano-4-aza-7,9-dideazaadenosine (Rem) in four steps (**Figure 2a**, **Methods, Supplementary Information**). The presence of RMP in the obtained RNA oligonucleotides was confirmed by denaturing HPLC and LC-MS after digestion into mononucleosides (**Figure 2b, c**). We further confirmed that the presence of RMP inhibits RNA extension by RdRp on minimal RNA template-product scaffolds (**Figure 2d, e**).

**Figure 2.**
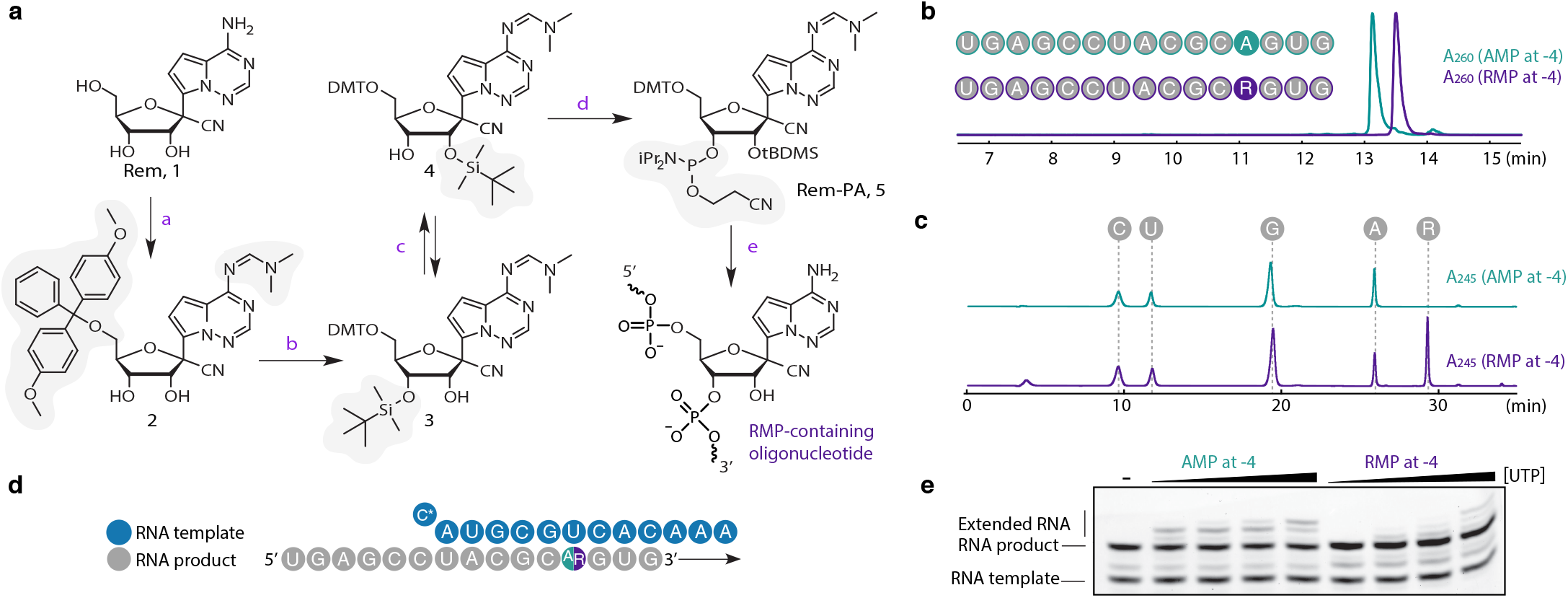
Preparation of remdesivir-containing RNA. **a** Scheme of the synthesis of 5’-O-DMT-2’-*O*-TBDMS-protected 3’-cyanoethyl diisopropyl phosphoramidite (Rem-PA), which was used to synthesize RNA oligos with remdesivir monophosphate (RMP) at defined positions. **b** Analysis of RMP-containing RNA by denaturing HPLC confirms the presence of RMP. **c** Analysis of the RMP-containing RNA by LC-MS after digestion into mononucleosides confirms the presence of RMP. **d** Minimal RNA template-product scaffold with RMP (R) or AMP (A) in a synthesized RNA oligonucleotide product strand. **e** The presence of RMP in a synthesized RNA oligonucleotide inhibits RNA extension by RdRp on the minimal RNA scaffold (**d**).

The ability to prepare RNAs containing RMP at defined positions enabled us to structurally trap the two states of the RdRp complex that are relevant for understanding remdesivir-induced RdRp stalling. Specifically, we investigated RdRp-RNA complexes trapped after addition of two or three nucleotides following RMP incorporation. We prepared RNA scaffolds containing RMP at positions –3 or –4 by annealing short RMP-containing oligonuclotides to long, loop-forming RNAs (scaffolds 1 and 2, respectively) (**Methods**). The annealed RNA scaffolds were then bound to purified RdRp and subjected to cryo-EM analysis as described^20^, resulting in two refined structures (**Extended Data Table 1**).

The first RdRp-RNA structure (structure 1) was resolved at 3.1 Å resolution (**Extended Data Figure 1**) and showed that the RMP was located at position –3 of the RNA product strand, as expected from the design of scaffold 1 (**Figure 3a, b**). The RdRp-RNA complex adopted the post-translocated state. The RNA 3’-end resided in the –1 site and the +1 site was free to bind the next NTP substrate. Comparison with our previous RdRp-RNA complex structure^20^ did not reveal significant differences. The 1’-cyano group of the RMP ribose moiety was located at position –3 and is accommodated there by an open space in the RNA product-binding site of the RdRp (**Figure 3a, b**). Thus structure 1 represents an active state of the elongation complex that is poised to add one more nucleotide to the RNA before stalling, consistent with biochemical results.

**Figure 3.**
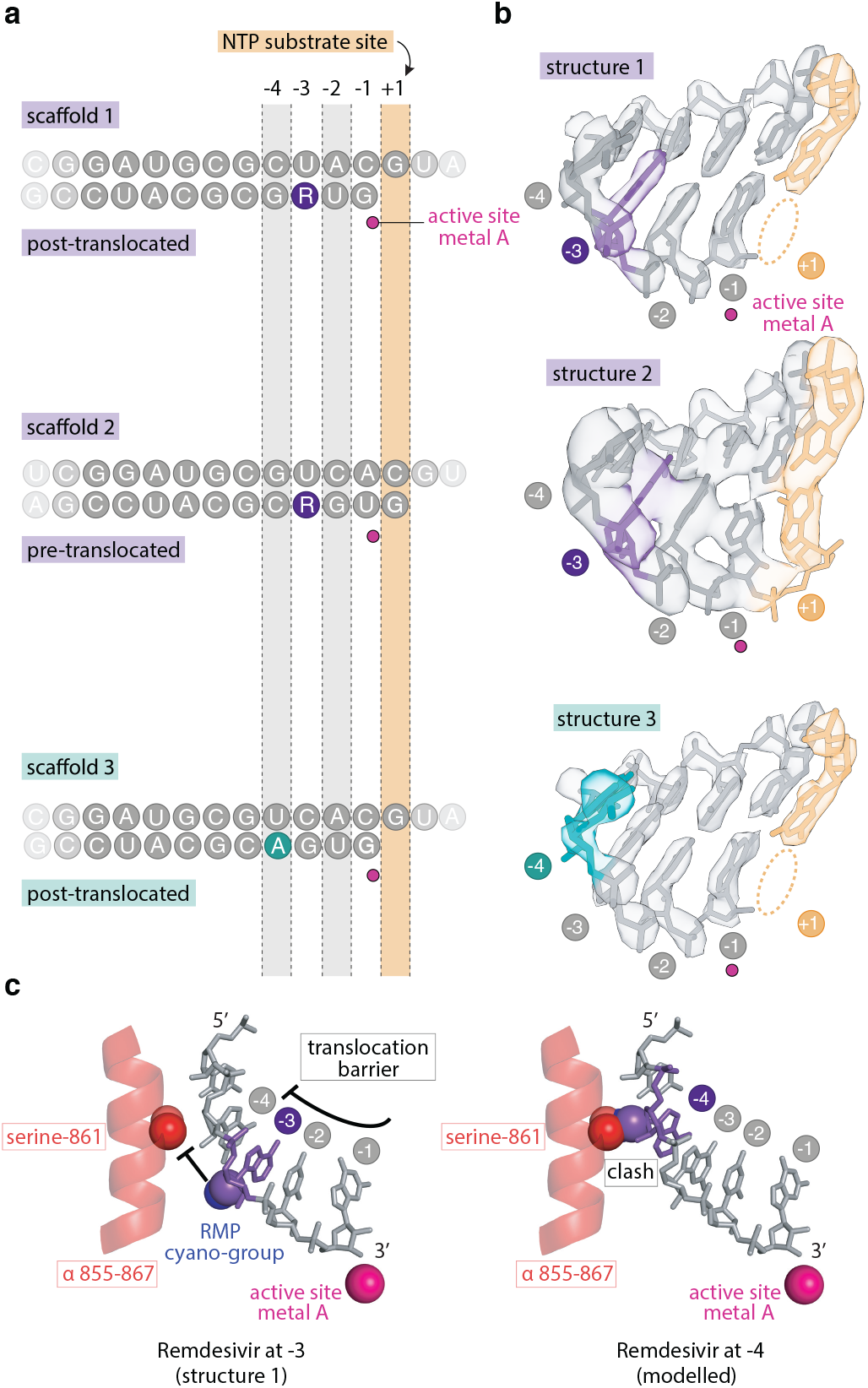
Structural analysis of remdesivir-induced RdRp stalling. **a** Position of RNA scaffolds 1–3 as observed in RdRp-RNA complex structures 1–3. Template and product strands are on the top and bottom, respectively. **b** Cryo-EM density of RNA in the active center of structures 1–3. The active site metal ion was modelled^30^ and is shown as a magenta sphere. **c** The C1’-cyano group of the RMP ribose moiety (violet) is accommodated at position –3 (left), but would clash with the side chain of nsp12 residue serine-861 (red) at position –4 (right). Spheres indicate atomic van der Waals surfaces.

The second RdRp-RNA structure (structure 2) was resolved at 3.4 Å resolution (**Extended Data Figure 1**) and showed that the RMP moiety was not located at position –4, as was expected from our design of scaffold 2, but was instead located at position –3 (**Figure 3a, b**). The +1 site was no longer free, as observed in structure 1, but was occupied by the nucleotide at the RNA 3’-end. The RdRp-RNA complex adopts the pre-translocated state and cannot bind the next NTP substrate. Thus structure 2 indicates that the RMP moiety in the RNA product strand is not tolerated at position –4. These results suggested that remdesivir-induced stalling of the RdRp is due to impaired translocation of the RNA after the RMP reaches register –3.

To test the hypothesis that RdRp stalling results from a translocation barrier, we formed a third RdRp-RNA complex with an RNA scaffold that was identical to that in structure 2 except that RMP was replaced by AMP, and we determined the resulting structure 3 at 2.8 Å resolution (**Figure 3a, b**, **Extended Data Figure 1**). In structure 3, the RdRp-RNA complex adopted the post-translocation state and the +1 site was again free, as observed in structure 1. This shows that the unexpected pre-translocated state that we observed in structure 2 was indeed caused by the presence of RMP, which was not tolerated at position –4. In conclusion, the RMP moiety in the RNA product strand gives rise to a translocation barrier that impairs movement of the RMP from position –3 to position –4.

Prior observations suggest that the translocation barrier that we observe here is caused by the presence of the C1’-cyano group in the remdesivir ribose moiety. First, this cyano group is critical for antiviral potency^7^. Second, modeling the RMP at position –4 of the RNA product strand results in a steric clash with the side chain of serine-861 in nsp12^6,8^. Indeed, our structural data strongly support the prior modeling (**Figure 3c**). Third, truncation of serine-861 to alanine^8,28^ or glycine^28^ renders the RdRp less sensitive or insensitive, respectively, to inhibition by remdesivir. We conclude that the translocation barrier that we observe here results from the sterically impaired passage of the cyano group in RMP past the serine-861 side chain in the nsp12 subunit of RdRp. We have summarized the mechanism of remdesivir-induced RdRp stalling in a molecular animation (**Supplementary Video**).

Finally, our results suggest how the remdesivir-stalled state escapes viral proofreading. In our structure 2, the RNA product 3’-nucleotide is matched and base-paired with the RNA template strand (**Figure 3a, b**). The matched RNA 3’-end may escape proofreading because the viral exonuclease nsp^14^ preferentially recognizes a mismatched 3’-end^9,10^. Nevertheless, some proofreading can occur and this renders remdesivir less efficient^5^, indicating that the viral exonuclease can remove several nucleotides from the base-paired RNA 3’-end. Such removal of several RNA nucleotides may require RNA backtracking along the RdRp, and this may be induced by the viral helicase nsp13^29^. While these issues will be studied further, our mechanistic in-sights may already facilitate the search for compounds with improved potential to interfere with coronavirus replication.

## EXPERIMENTAL PROCEDURES

No statistical methods were used to predetermine sample size. The experiments were not randomized, and the investigators were not blinded to allocation during experiments and outcome assessment.

### RNA extension assays

Preparation of SARS-CoV-2 RdRp was carried out as described^20^. RTP synthesis is described in Supplementary Information. All unmodified RNA oligonucleotides were purchased from Integrated DNA Technologies. The RNA sequence used for the RNA extension assay (Figure 1c) is /56-FAM/rArArC rArGrG rArGrA rCrUrC rGrCrG rUrArG rUrUrUrU rCrUrA rCrGrC rG. The assay was performed as described^20^, except for the following changes. The final concentrations of nsp12, nsp8, nsp7 and RNA were 3 μM, 9 μM, 9 μM and 2 μM, respectively. The highest concentration of NTPs was 0.5 mM for each nucleotide (ATP or RTP, GTP, UTP and CTP), followed by a two-fold serial dilution. RNA products were resolved on a denaturing sequencing gel and visualized by Typhoon 95000 FLA Imager (GE Healthcare Life Sciences). The RNA sequences used for extending RMP-containing RNA oligonucleotides are: rUrGrA rGrCrC rUrArC rGrC-rA or rR-rGrUrG (product) and rAr-ArA rCrArC rUrGrC rGrUrA /3ddC/ (template). The extension assay (Figure 2e) was performed as described^20^, with minor changes. Reactions were started by addition of UTP (final concentrations: 6.25 μM, 12.5 μM, 25 μM or 250 μM). RNA products were visualized by SYBR Gold (Thermo Fischer) staining and imaged with Typhoon 9500 FLA Imager (GE Healthcare Life Sciences).

### Preparation of RMP-containing RNA oligonucleotides

1’-cyano-4-aza-7,9-dideazaadenosine (Rem, **1**) was converted to the 5’-*O*-DMT-2’-*O*-TBDMS-protected 3’-cyanoethyl diisopropyl phosphoramidite (Rem-PA, **5**) in four steps. Details of the synthetic procedures and NMR spectra of isolated compounds are given in the Supplementary Information. RNA oligonucleotides were then prepared by solid-phase synthesis on CPG support (0.6 μmol scale) using 2’-*O*-TOM-protected ribonucleoside phosphoramidites (100 mM in CH_3_CN) and ethylthiotetrazol (ETT, 250 mM in CH_3_CN) as activator, with 4 min coupling time. Rem-PA was used as freshly prepared solution (100 mM) in dry 1,2-dichloroethane, with a coupling time of two times 12 min. Detritylation was performed with 3% dichloroacetic acid in dichloromethane. Capping solutions contained 4-dimethylamino pyridine (0.5 M) in acetonitrile for Cap A, and acetic anhydride/sym-collidine/acetonitrile (2/3/5) for Cap B. Oxidation was performed with iodine (10 mM) in acetonitrile/sym-collidin/water (10/1/5). The oligonucleotides were deprotected with 25% NH_4_OH at 25 °C for 30 h, followed by 1 M TBAF in THF for 12 h, and purified by denaturing poly-acrylamide gel electrophoresis.

### Analysis of RMP-containing RNA oligonucleotides

The purity and identity of the RNA oligonucleotides was analyzed by anion-exchange HPLC (Dionex DNAPac PA200, 2×250 mm, at 60 °C. Solvent A: 25 mM Tris-HCl (pH 8.0), 6 M Urea. Solvent B: 25 mM Tris-HCl (pH 8.0), 6 M Urea, 0.5 M NaClO_4_. Gradient: linear, 0–40% solvent B, 4% solvent B per 1 CV), and HR-ESI-MS (Bruker micrOTOF-Q III, negative ion mode, direct injection). An aliquot (200 pmol in 25 μL) was digested by snake venom phosphodiesterase (SVPD, 0.5 U) in the presence of bacterial alkaline phosphatase (BAP, 0.5 U) in 40 mM Tris. pH 7.5, 20 mM MgCl_2_, and the resulting mononucleosides were analyzed by LC-ESI-MS using an RP-18 column (Synergi 4 μm Fusion-RP C18 80 Å, 250 x 2 mm, at 25 °C. aqueous mobile phase A: 5 mM NH_4_OAc, pH 5.3. organic mobile phase B: 100% acetonitrile. Gradient: 0–5% B in 15 min, then 5–50% B in 20 min, flow rate 0.2 mL/min) and micrOTOF-Q III with ESI ion source operated in positive ion mode (capillary voltage: 4.5 kV, end plate offset: 500 V, nitrogen nebulizer pressure: 1.4 bar, dry gas flow: 9 L/min). Extracted ion chromatograms and UV absorbance traces at 245 nm confirmed presence of remdesivir.

### Cryo-EM sample preparation and data collection

SARS-CoV-2 RdRp was prepared as described^20^. RNA scaffolds for RdRp-RNA complex formation were prepared by mixing equimolar amounts of two RNA strands in annealing buffer (10 mM Na-HEPES pH 7.4, 50 mM NaCl) and heating to 75 °C, followed by step-wise cooling to 4 °C. RNA sequences for RMP at position −3 (structure 1) are: rUrUrU rUrCrA rUrGrC rArU-rC rGrCrG rUrArG rGrCrU rCrArU rArCrC rGrUrA rUrUrG rArGrA rCrCrU rUrUrU rGrGrU rCrUrC rArArU rArCrG rGrUrA and rUrGrA rGrCrC rUrArC rGrCrG rRrUrG. RNA sequences for AMP/RMP at position −4 (structures 2 and 3) are: rUrUrU rUrCrA rUrGrC rArCrU rGrCrG rUrArG rGrCrU rCrArU rArCrC rGrUrA rUrUrG rArGrA rCrCrU rUrUrU rGrGrU rCrUrC rArArU rArCrG rGrUrA and rUrGrA rGr-CrC rUrArC rGrC- rA/rR -rGrUrG. RdRp-RNA complexes were formed by mixing purified nsp12 (scaffold 1: 2 nmol, scaffold 2: 2 nmol, scaffold 3: 1.6 nmol) with an equimolar amount of annealed RNA scaffold and 3-fold molar excess of each nsp8 and nsp7. After 10 minutes of incubation at room temperature, the mixture was applied to a Superdex 200 Increase 3.2/300 size exclusion chromatography column, which was equilibrated in complex buffer (20 mM Na-HEPES pH 7.4, 100 mM NaCl, 1 mM MgCl_2_, 1 mM TCEP) at 4 °C. Peak fractions corresponding to the RdRp-RNA complex were pooled and diluted to approximately 2 mg/ml. For structure 2, an additional 0.36 nmol of annealed RNA scaffold were spiked into the sample prior to grid preparation. 3 μL of the concentrated RdRp-RNA complex were mixed with 0.5 μl of octyl ß-D-glucopyranoside (0.003% final concentration) and applied to freshly glow discharged R 2/1 holey carbon grids (Quantifoil). The grids were blotted for 5 seconds using a Vitrobot MarkIV (Thermo Fischer Scientific) at 4 °C and 100% humidity and plunge frozen in liquid ethane.

Cryo-EM data were collected using SerialEM^31^ on a 300 keV Titan Krios transmission electron microscope (Thermo Fischer Scientific). Prior to detection, inelastically scattered electrons were filtered out with a GIF Quantum energy filter (Gatan) using a slit width of 20 eV. Images were acquired in counting mode (non-super resolution) on a K3 direct electron detector (Gatan) at a nominal magnification of 105,000x resulting in a calibrated pixel size of 0.834 Å/pixel. Images were exposed for a total of 2.2 seconds with a dose rate of 19 e^−^/px/s resulting in a total dose of 60 e^−^/Å^2^, which was fractionated into 80 frames. Previous cryo-EM analysis of the SARS-CoV2 RdRp showed strong preferred orientation of the RdRp particles in ice^20^.

Therefore, all data were collected with 30° tilt to obtain more particle orientations. Motion correction, CTF-estimation, and particle picking and extraction were performed on the fly using Warp^32^. In total, 8,004, 11,764 and 7,043 movies were collected for structures 1, 2 and 3, respectively.

### Cryo-EM data processing and analysis

For structure 1, 1.8 million particles were exported from Warp^32^ 1.0.9 to cryoSPARC^33^ 2.15. After *ab initio* refinement of five classes, an intermediate map from the previous processing of EMD-11007^20^ was added as a 6^th^ reference, and supervised 3D classification was performed. 654k particles (37%) from the best class deemed to represent the polymerase were subjected to 3D refinement to obtain a 3.1 Å map. Half-maps and particle alignments were exported to M^34^ 1.0.9, where reference-based frame series alignment with a 2×2 image-warp grid as well as CTF refinement were performed for two iterations. Although the resulting map had the same 3.1 Å nominal resolution, the features of interest were significantly cleaner.

For structure 2, 3.4 million particles were exported from Warp 1.0.9 to cryoSPARC 2.15. After *ab initio* refinement of 5 classes, the EMD-11007-related reference was added as a 6^th^ reference, and supervised 3D classification was performed. To further clean up the resulting 881k particles (26%) from the best class deemed to represent the polymerase, another *ab initio* refinement of five classes was performed. Three of these classes and the EMD-11007-related reference were used for supervised 3D classification. 474k particles (54%) from the best class deemed to represent the polymerase were subjected to 3D refinement to obtain a 3.6 Å map. Half-maps and particle alignments were exported to M 1.0.9, where reference-based frame series alignment with a 2×2 image-warp grid as well as CTF refinement were performed for two iterations to obtain a 3.4 Å map.

For structure 3, 2.2 million particles were exported from Warp 1.0.9 to cryoSPARC 2.15. Initial unsupervised 3D classification in three classes was performed using the EMD-11007-related reference. To further clean up the resulting 1.1 million particles (23%), *ab initio* refinement of 5 classes was performed. Four of these classes and the EMD-11007-related reference were used for supervised 3D classification. 819k particles (70%) from the best class deemed to represent the polymerase were subjected to 3D refinement to obtain a 3.1 Å map. Half-maps and particle alignments were exported to M 1.0.9, where reference-based frame series alignment with a 2×2 image-warp grid as well as CTF refinement were performed for three iterations to obtain a 2.8 Å map.

### Model building and refinement

Models were built using our previously published SARS-CoV-2 RdRp structure as starting model (PDB 6YYT)^20^. For each of the structures 1–3, the model was first rigid-body fit into the density and subsequently adjusted in real-space in Coot^35^. Parts of the N-terminal NiRAN domain of nsp12, the N-terminal extension of nsp8a and the entire nsp8b molecule were removed due to absence or poor quality of density for these regions. Restraints for RMP were generated in phenix.elbow^36^ and the structures were refined using phenix.real_space_refine^37^ with appropriate secondary structure restraints. Model quality was assessed using MolProbity within Phenix^38^ which revealed excellent stereochemistry for all three structural models (Extended Data Table 1). Figures were prepared with PyMol and ChimeraX^39^.

## ACKNOWLEDGEMENTS

We thank Jan Seikowski, Jürgen Bienert and Vladimir Belov for synthesis of RTP. H.S.H. was supported by the Deutsche Forschungsgemeinschaft (FOR2848). C.H. was supported by the DFG (SPP1784) and the ERC Consolidator Grant illumizymes (grant agreement No 682586). P.C. was supported by the Deutsche Forschungsgemeinschaft (SFB860, SPP2191, EXC 2067/1-390729940) and the ERC Advanced Investigator Grant CHROMATRANS (grant agreement No 693023).

## AUTHOR CONTRIBUTIONS

G.K., H.S.H., D.T., and C.D. designed and carried out biochemical and structural biology experiments and analyzed corresponding data. F.S. and A.S. synthesized and analyzed RMP-containing RNA oligonucleotides. J.S. and L.F. prepared RdRp. C.H. designed and supervised synthesis of RMP-containing RNA. P.C. designed and supervised research. P.C. wrote the manuscript, with input from all authors.

## COMPETING INTERESTS

The authors declare no competing interests.

## DATA AVAILABILITY

Reconstructions and structure coordinate files will be made available in the Electron Microscopy Database and the Protein Data Bank.

## SUPPLEMENTARY MATERIALS

**Supplementary Information | Synthesis of RMP-containing RNA oligonucleotides. Related to** **Figures 1, 2.**

**Supplementary Video | Animation of remdesivir-induced RdRp stalling.**

Remdesivir (purple, C1’-cyano group in van der Waals surface rendering) binds to the +1 site and base pairs with a uridine in the template strand (blue) and is then incorporated into the growing RNA product (light red). For simplicity, NTP binding and release of pyrophosphate were omitted and instead only the RMP moiety is shown as it is added to RNA. Translocation of the RNA then transfers the RNA product 3’-nucleotide from position +1 to position –1. Such nucleotide addition cycle is repeated three more times using cognate, natural nucleotides (red). After addition of the third nucleotide, translocation is impaired because the C1’-cyano group of remdesivir encounters the side chain of serine-861 (shown in van der Waals surface rendering) of RdRp subunit nsp12. The translocation barrier is indicated by failed attempts to translocation that are shown twice for clarity. Location of the RdRp active site is indicated by a magenta sphere.

---

**Extended Data Figure 1.**
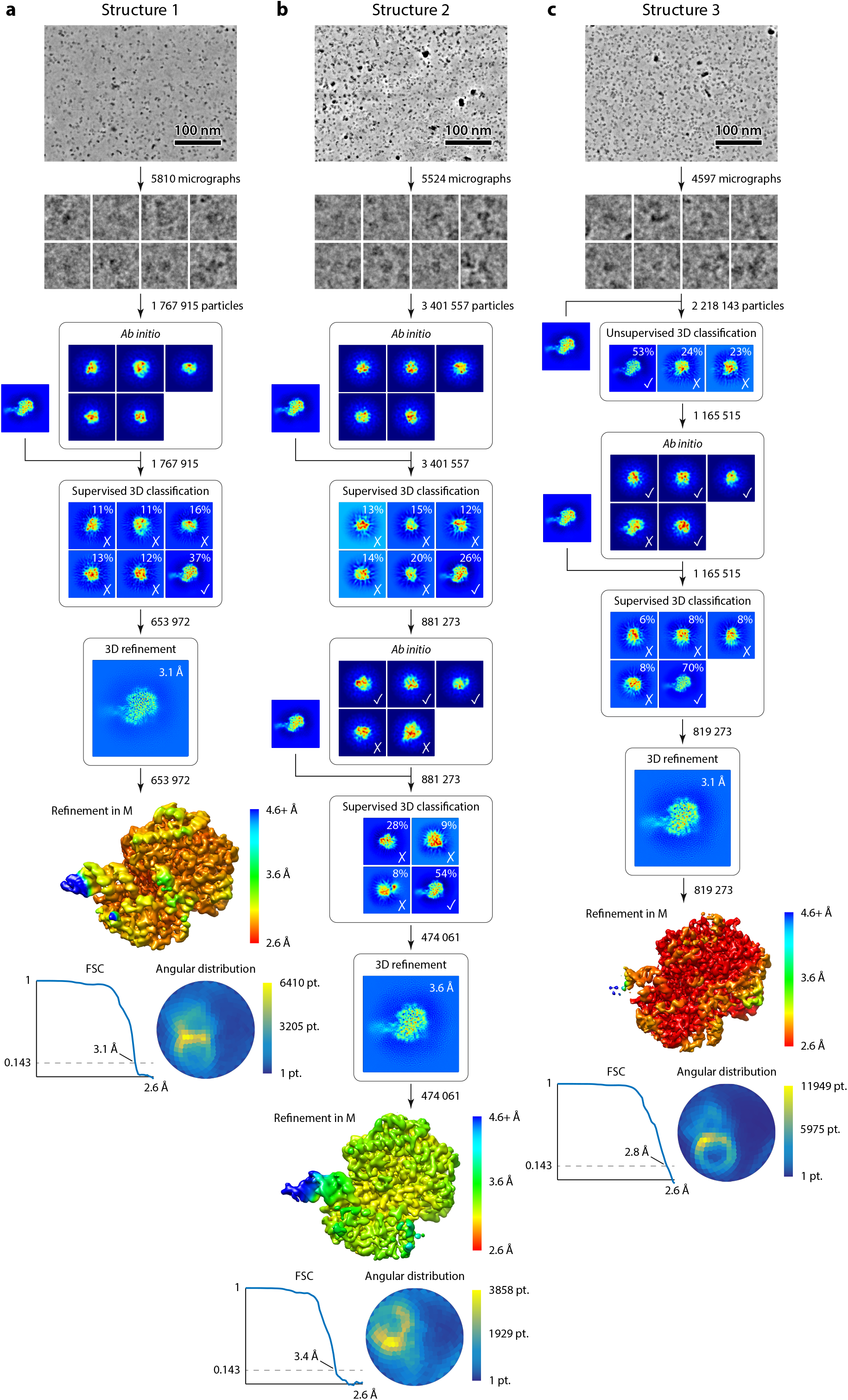
Cryo-EM sorting trees and quality of reconstructions. Related to Figure 3. Cryo-EM sorting tree (top); local resolution, FSC plot and angular distribution of the final reconstruction (bottom) for RMP-containing RdRp-RNA structure 1 (**a**), RMP-containing RdRp-RNA structure 2 (**b**), and RdRp-RNA structure 3 (**c**).

## EXTENDED DATA TABLE

**Extended Data Table 1.**
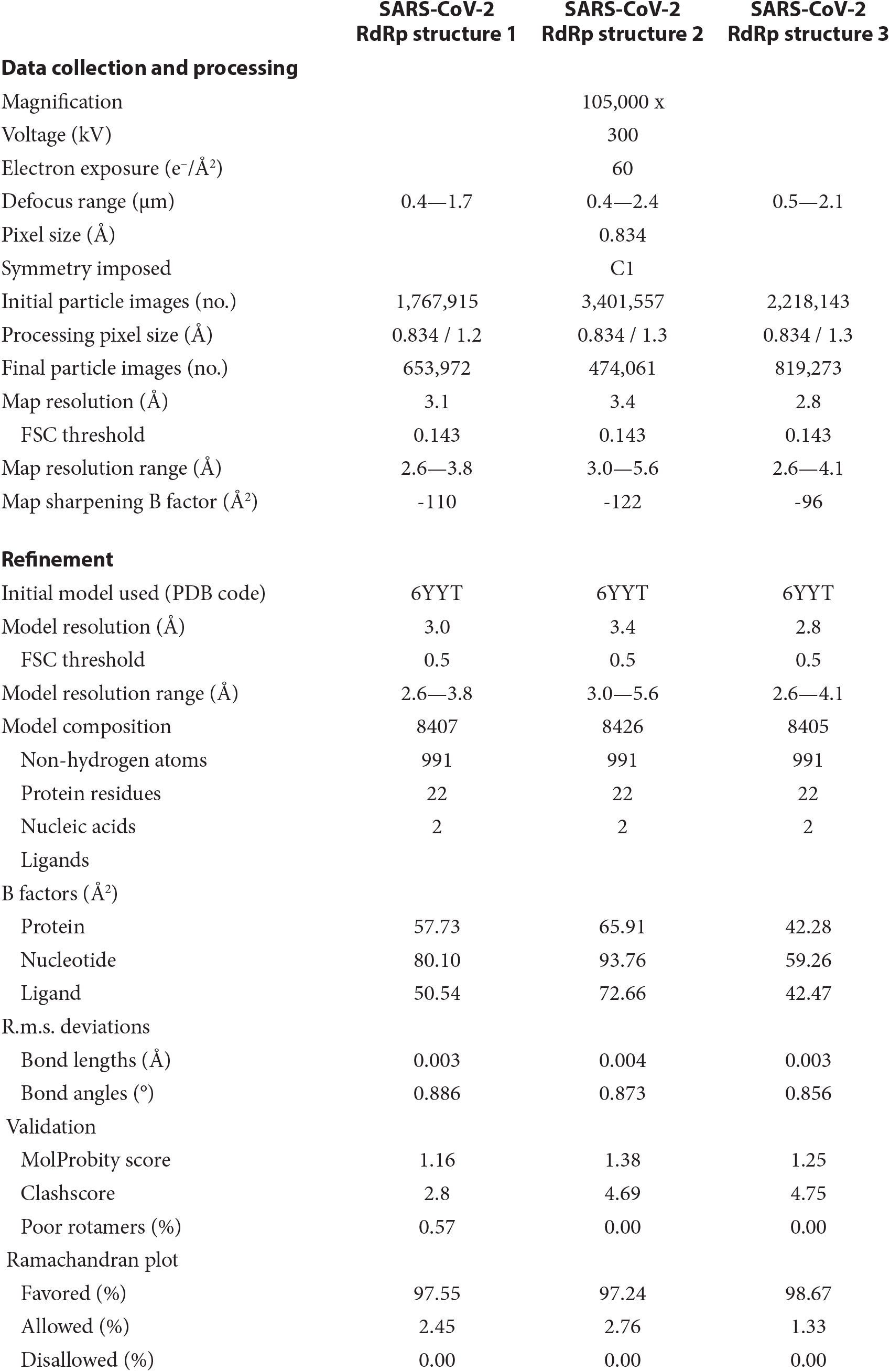
Cryo-EM data collection, refinement, and validation statistics.

|  | SARS-CoV-2<br>RdRp structure 1 | SARS-CoV-2<br>RdRp structure 2 | SARS-CoV-2<br>RdRp structure 3 |
| --- | --- | --- | --- |
| **Data collection and processing** |  |  |  |
| Magnification |  | 105,000 x |  |
| Voltage (kV) |  | 300 |  |
| Electron exposure (e<sup>-</sup>/Å<sup>2</sup>) |  | 60 |  |
| Defocus range (μm) | 0.4—1.7 | 0.4—2.4 | 0.5—2.1 |
| Pixel size (Å) |  | 0.834 |  |
| Symmetry imposed |  | C1 |  |
| Initial particle images (no.) | 1,767,915 | 3,401,557 | 2,218,143 |
| Processing pixel size (Å) | 0.834 / 1.2 | 0.834 / 1.3 | 0.834 / 1.3 |
| Final particle images (no.) | 653,972 | 474,061 | 819,273 |
| Map resolution (Å) | 3.1 | 3.4 | 2.8 |
| FSC threshold | 0.143 | 0.143 | 0.143 |
| Map resolution range (Å) | 2.6—3.8 | 3.0—5.6 | 2.6—4.1 |
| Map sharpening B factor (Å<sup>2</sup>) | -110 | -122 | -96 |
| **Refinement** |  |  |  |
| Initial model used (PDB code) | 6YYT | 6YYT | 6YYT |
| Model resolution (Å) | 3.0 | 3.4 | 2.8 |
| FSC threshold | 0.5 | 0.5 | 0.5 |
| Model resolution range (Å) | 2.6—3.8 | 3.0—5.6 | 2.6—4.1 |
| Model composition | 8407 | 8426 | 8405 |
| Non-hydrogen atoms | 991 | 991 | 991 |
| Protein residues | 22 | 22 | 22 |
| Nucleic acids | 2 | 2 | 2 |
| Ligands |  |  |  |
| B factors (Å<sup>2</sup>) |  |  |  |
| Protein | 57.73 | 65.91 | 42.28 |
| Nucleotide | 80.10 | 93.76 | 59.26 |
| Ligand | 50.54 | 72.66 | 42.47 |
| R.m.s. deviations |  |  |  |
| Bond lengths (Å) | 0.003 | 0.004 | 0.003 |
| Bond angles (°) | 0.886 | 0.873 | 0.856 |
| Validation |  |  |  |
| MolProbity score | 1.16 | 1.38 | 1.25 |
| Clashscore | 2.8 | 4.69 | 4.75 |
| Poor rotamers (%) | 0.57 | 0.00 | 0.00 |
| Ramachandran plot |  |  |  |
| Favored (%) | 97.55 | 97.24 | 98.67 |
| Allowed (%) | 2.45 | 2.76 | 1.33 |
| Disallowed (%) | 0.00 | 0.00 | 0.00 |

## Notes

### Competing Interest Statement

The authors have declared no competing interest.

