## Supplementary Information for "Mechanism of SARS-CoV-2 polymerase inhibition by remdesivir"

### Table of Contents

**General information** .......... 2
**Experimental procedures for synthesis of RTP (to Figure 1)** .......... 2-5
**Experimental procedures for synthesis of Rem-PA (to Figure 2)** .......... 6-11
**NMR Spectra** .......... 12-20
References .......... 21

#### Supplementary Figures

### General information

All reactions were performed under inert nitrogen atmosphere with dry solvents. Reagents used for synthesis were purchased in 'pro analysis' or 'pro synthesis' quality and used without further purification. Solvents used for synthesis were purchased in 'puriss. over molecular sieves', 'pro analysis' or 'pro synthesis' quality and used without further purification. For column chromatography, solvents in technical quality were purchased and purified by distillation. For solid-phase synthesis, acetonitrile and dichloroethane were dried over molecular thieves. Thin layer chromatography (TLC) was performed on aluminum plates pre-coated with silica gel 60 $F_{254}$ (Merck). Substances were detected based on fluorescence quenching at 254 nm. For column chromatography, silica gel 60 (Merck) with a particle size of 40 – 63 µm was used. NMR spectra were recorded using Bruker Avance III (400 MHz) spectrometers. Chemical shifts ($\delta$) are given in ppm and were referenced using the deuterated solvent as internal standard. Data are reported as: s = singlet, d = doublet, t = triplet, q = quartet, m = multiplet, br = broad; Coupling constants ($J$) are given in Hz. High-resolution (HR) electrospray ionization (ESI) mass spectra (MS) were recorded on a Bruker micrOTOF-Q III spectrometer. The detected mass-to-charge ratio ($m/z$) is given, as well as the calculated monoisotopic mass.

### Experimental procedures and compound characterization for RTP (to Figure 1)

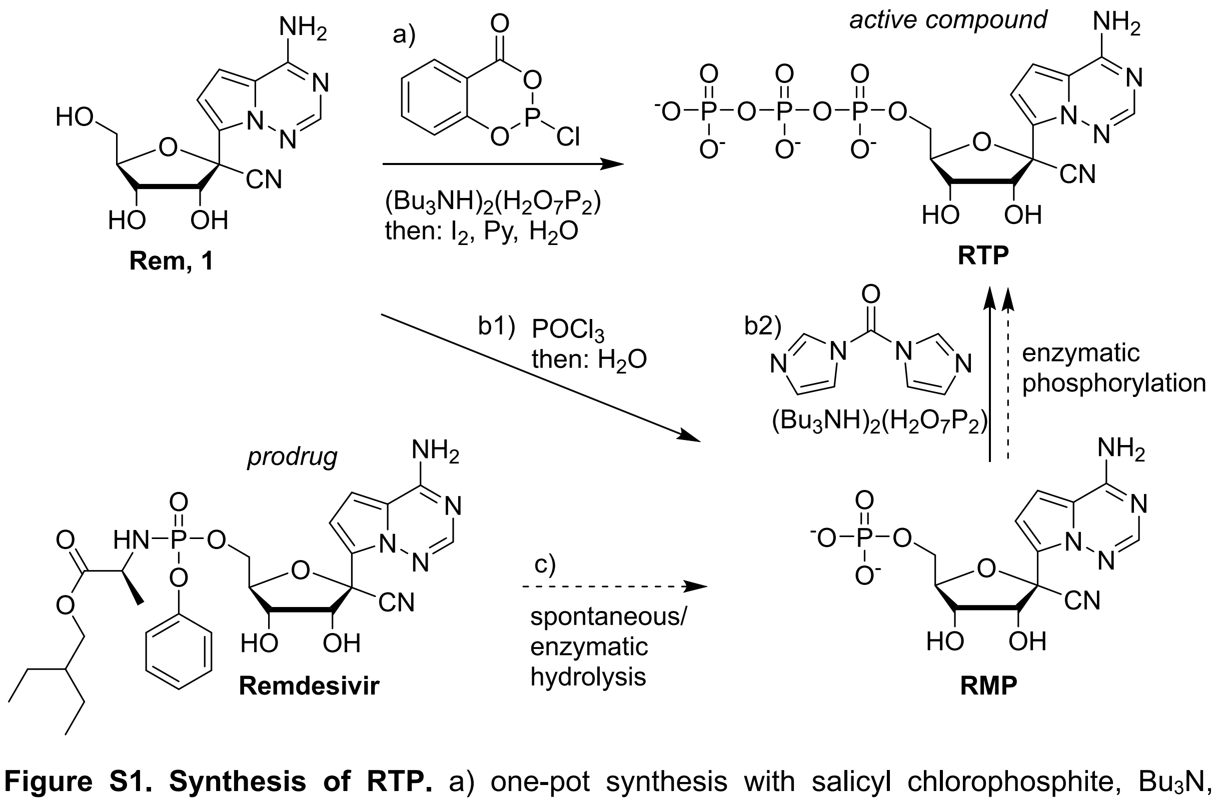

**Figure S1. Synthesis of RTP.** a) one-pot synthesis with salicyl chlorophosphite, $Bu_3N$, bis(tributylammonium)pyrophosphate in DMF followed by oxidation with iodine in aqueous pyridine.[1] b) two-step synthesis with isolation of RMP. b1) $POCl_3$, $PO(OMe)_3$, then $Et_3NHCO_3$ buffer. b2) carbonyldiimidazole, bis(tributylammonium)pyrophosphate in DMF. c) enzymatic (intracellular) activation of Remdesivir for comparison.

1'-Cyano-4-aza-7,9-dideazaadenosine 5'-triphosphate (**RTP**) triethylammonium salt

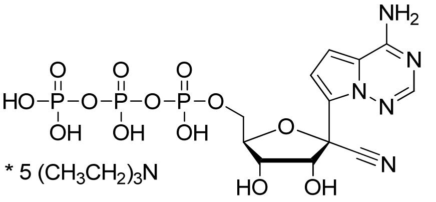

**RTP**
$C_{12}H_{16}N_5O_{13}P_3$
531.20 g•mol$^{-1}$

a) one-pot procedure following a general method described by Caton-Williams et al.[1]

Bis(tributylammonium)pyrophosphate (154 mg, 280 µmol, 2.00 eq) was dissolved in dry DMF (0.48 mL), tri-*n*-butylamine (520 µL, 2.2 mmol, 16 eq) was added, and the reaction mixture was stirred at ambient temperature for 5 min. A solution of salicyl chlorophosphite (57 mg, 280 µmol, 2.00 eq) in anhydrous DMF (0.48 mL) was added, and the reaction mixture was stirred vigorously at ambient temperature for 30 min.

Two equivalents of the in-situ generated triphosphorylation reagent were added at 0°C to 1'-cyano-4-aza-7,9-dideazadenosine (**1**, 19.5 mg, 67 µmol, 1.00 eq.). After removal of the ice bath, the reaction mixture was stirred for 3 h at ambient temperature. A solution of iodine (20 mM in pyridine/water 9:1, ca. 1.1 mL) was added stepwise, until a brown color persisted for at least 15 min. Two reaction volumes of ultrapure water (ca. 3.3 mL) were added, followed by stirring for 1.5 h at ambient temperature. An aqueous solution of sodium chloride (20% in water), followed by ethanol (ca. 22 mL) were added, and the reaction mixture kept on dry ice for 1 h. The crude mixture was centrifuged for 10 min at -9°C (3200g), the supernatant removed, and the air dried pellet was purified by RP HPLC using a gradient of 3 % to 10 % acetonitrile in triethylammonium acetate buffer (50 mM, pH 7.5). The purest fractions were pooled, the solvent removed by lyophilization, and the product dissolved in water (350 µL) to yield a stock solution of **RTP.** The concentration was determined by UV spectroscopy on a NanoDrop One spectrometer (Thermo Fisher Scientific) using $\varepsilon^{245}$ = 37350 M$^{-1}$cm$^{-1}$ to give a concentration of 10 mM (3.5 µmol, 5.2 % yield).

**$^1$H NMR** (400 MHz, $D_2O$): δ (ppm) = 7.92 (s, 1H, C2-H), 7.08 (d, *J* = 4.8 Hz ,1H, C5-H), 6.94 (d, *J* = 4.7 Hz ,1H, C6-H), 5.02 (d, *J* = 5.3 Hz, 1H, C2'-H), 4.60 (dd, *J* = 5.3 and 3.1 Hz, 1H, C4'-H), 4.51 (m, 1H, C3'-H), 4.21 (m, 1H, C5'-H), 4.01 (ddd, 1H, *J* = 11.8, 4.7 and 3.1 Hz, C5'-H), 3.15 (q, ~30H, $CH_2N$ in triethylamine), 1.23 (t, ~45H, $CH_3$ in triethylamine).

**$^{13}$C (gHSQCAD) NMR** (101 MHz, $D_2O$): δ (ppm) = 147.30 (C2), 111.3 (C5), 102.4(C6), 85.5 (C3'), 74.9 (C2'), 70.1 (C4'), 64.7 (C5'), resonances of the protonated carbon atoms were recorded.

**$^{31}$P{$^1$H} NMR** (394 MHz, $D_2O$): δ (ppm) = -6.5 (d, *J* = 20 Hz), -11. 4 (d, *J* = 20 Hz ),-22.5 (t, *J* = 20 Hz).

**HR-ESI-MS**: *m*/*z* calc. ($C_{12}H_{15}N_5O_{13}P_3$ [M-H]$^-$): 529.98792, found: 529.98842.

Analytical data are consistent with previously reported values.[2]

b) Two-step procedure with isolation of RMP

1'-Cyano-4-aza-7,9-dideazaadenosine 5'-monophosphate (**RMP**)

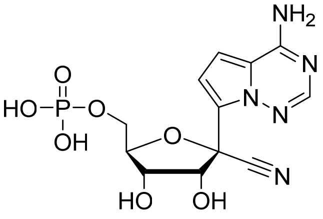

**RMP**
$C_{12}H_{14}N_5O_7P$
371.25 g•mol$^{-1}$

1'-cyano-4-aza-7,9-dideazadenosine (**1**, 51 mg, 0.17 mmol, 1.00 eq.) was dissolved in dry trimethylphosphate (2.5 mL), cooled to 0 °C, treated with $POCl_3$ (0.3 mL, 0.34 mmol, 1.90 eq.) and stirred at 0 °C for 4 h. A solution of tributylamine (0.2 mL, 0.84 mmol, 4.9 eq.) and bis(tributylammonium)pyrophosphate (150 mg, 0.27 mmol, 1.6 eq.) in dry MeCN (5 mL) was added and the mixture stirred at 0 °C for 0.5 h.) Triethylammonium bicarbonate (TEAB) buffer (1 M, pH 7.5, 2 mL) was added at 0 °C and the mixture stirred at ambient temperature for 0.5 h. The solvent was removed under reduced pressure and coevaporated with water. The crude product was subjected to ion exchange chromatography using a gradient of 0 % to 100 % TEAB buffer (1 M, pH 7.5) in water. After evaporation, the residue was further purified by RP HPLC using a gradient of 0 % to 40 % acetonitrile in triethylammonium acetate buffer (100 mM, pH 7.0). The solvent was removed in high vacuum to yield **RMP** as the triethylammonium salt (37 mg, 0.08 mmol, 46%). In contrast to a previous report,[2] no triphosphate was obtained, most likely because the pyrophosphate solution contained traces of water or the reaction time was too short. The obtained monophosphate was used as a reference compound for analysis of oligonucleotide digestion experiments and determination of the extinction coefficient (see UV spectrum in Figure S2b). The triphosphate (RTP) was obtained upon activation of the monophosphate in the next step.

**$^1$H NMR** (400 MHz, $D_2O$): δ (ppm) = 7.78 (s, 1H, C2-H), 6.84 (d, *J* = 4.7 Hz ,1H, C5-H), 6.67 (d, *J* = 4.7 Hz ,1H, C6-H), 4.86 (d, *J* = 5.5 Hz, 1H, C2'-H), 4.46 - 4.44 (m, 1H, C4'-H), 4.39 (dd, *J* = 4.3, 4.0 Hz, 1H, C3'-H), 4.02 - 3.99 (m, 2H, C5'-H).

**$^{13}$C{$^1$H} NMR** (125 MHz, $D_2O$): δ (ppm) = 154.30 (1C, C6), 145.87 (1C, C2), 123.20 (1C, C7), 116.88 (1C, CN), 115.80 (1C, C5), 111.10 (1C, C9), 102.72 (1C, C8), 84.95 (1C, C4'), 76.51 (1C, C1'), 74.68 (1C, C2'), 70.17 (1C, C3'), 63.99 (1C, C5').

**$^{31}$P{$^1$H} NMR** (202 MHz, $D_2O$): δ (ppm) = 0.47 (s).

**HR-ESI-MS**: *m*/*z* calc. ($C_{12}H_{13}N_5O_7P$ [M-H]$^-$): 370.05581, found: 370.05678.

1'-Cyano-4-aza-7,9-dideazaadenosine 5'-triphosphate (**RTP**)

The **RMP** triethylammonium salt (20 mg, 42 µmol, 1.00 eq.) was dissolved in dry DMF (1 mL), a solution of 1,1'-carbonyldiimidazole (25 mg, 151 µmol, 3.6 eq.) in dry DMF (0.5 mL) was added and the mixture stirred at ambient temperature for 18 h. Methanol (4.9 µL) was added and stirred at ambient temperature for 1 h. A solution of bis(tributylammonium)pyrophosphate (93 mg, 170 µmol, 4.1 eq.) in dry DMF (1 mL) was added and the mixture stirred at ambient temperature for 22 h. The solution was cleared from the precipitate, and the solvent was removed under reduced pressure. The residue was dissolved in water, washed with dichloromethane, and then water was removed under reduced pressure. The crude product was purified by ion exchange HPLC using a gradient of 0 % to 100 % TEAB buffer (1 M, pH 7.5) in water. The solvent was removed in high vacuum and the product dissolved in water to yield a solution of **RTP.** The concentration was determined by UV spectroscopy on a Cary 100 Bio spectrometer using $\varepsilon^{245} = 37350\ \text{M}^{-1}\text{cm}^{-1}$ to give a yield of (16 mM, 800 µL, 12.7 µmol, 30 %). The spectroscopic data were consistent with the ones reported in a) (see above). In addition, RTP and RMP were analyzed by anion exchange HPLC.

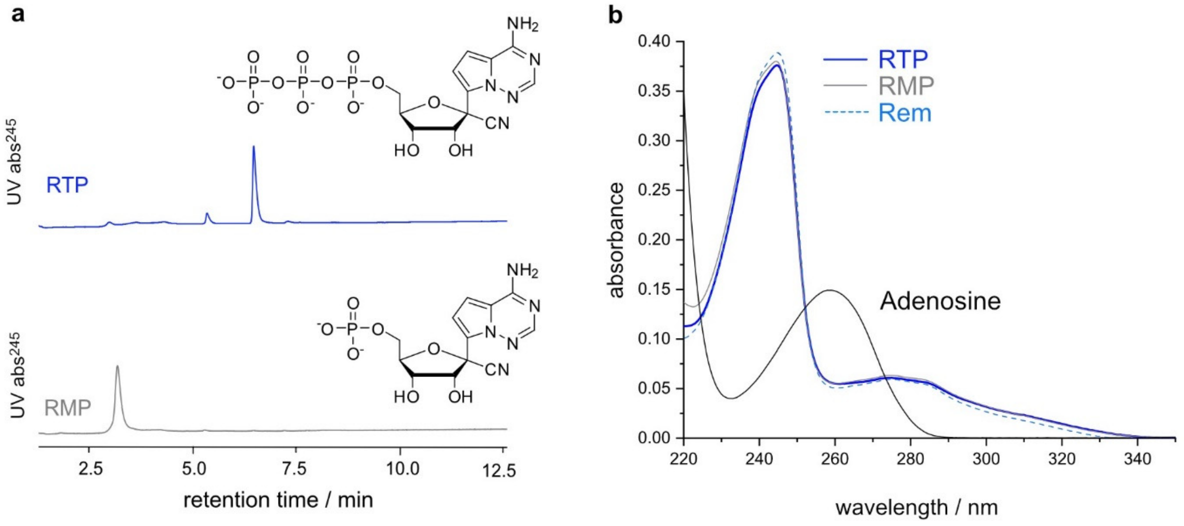

**Figure S2. Characterization of synthetic RMP and RTP**. a) Anion exchange HPLC on Dionex DNA Pac PA200, elution with linear gradient of sodium perchlorate in Tris.HCl buffer (25 mM, pH 8.0); UV Absorbance monitored at 245 nm. b) UV absorbance spectra of solutions of RTP, RMP, and Rem, in comparison to adenosine, in 10 mM Na phosphate buffer pH 7.4. c = 10 µM, d = 1 cm.

**Experimental procedures and compound characterization of Rem-PA (to Figure 2)**

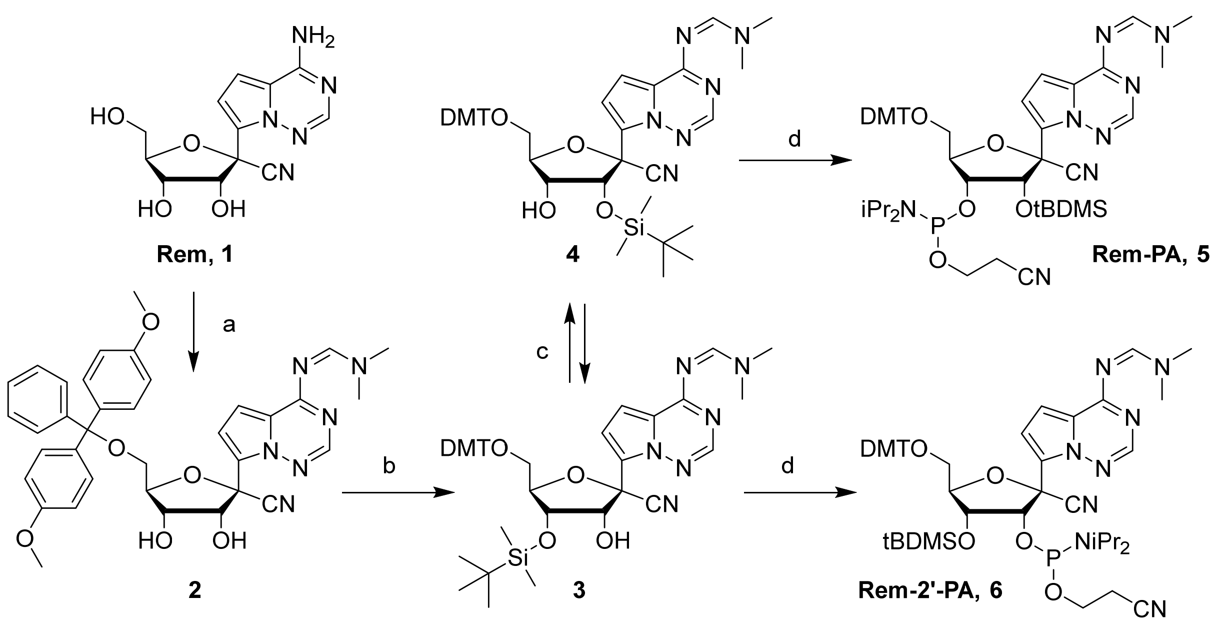

**Figure S3. Synthesis of Rem-PA.** a) i) DMFDMA, pyridine, rt, 18 h, ii) DMT-Cl, pyridine, rt, 3.5 h, iii) MeOH, rt, 30 min, 93%; b) tBDMS-Cl, $AgNO_3$, pyridine, rt, 22 h, 72%; c) MeOH, $Et_3N$, rt, 30 min, 30% (**4**); d) CEPCl, $EtNMe_2$, DCM, rt, 5 h, 75% (**5**), 85% (**6**).

1'-Cyano-4-aza-7,9-dideazaadenosine (Rem, **1**) was reacted with dimethylformamide dimethylacetal (DMFDMA) in pyridine, resulting in protection of $N^6$ and temporary 2',3'-acetal formation. Installation of the 5'-*O*-4,4'-dimethoxytrityl group was followed by release of the 2',3'-acetal and isolation of compound **2**. Treatment with tBDMS-Cl in the presence of $AgNO_3$ produced predominantly the 3'-*O*-tBDMS-protected nucleoside **3**, which was equilibrated in MeOH/$NEt_3$ to allow isolation of the 2'-*O*-tBDMS-protected isomer **4**. Both compounds **3** and **4** were individually converted to the corresponding 2-cyanoethyl diisopropylphosphoramidites **5** and **6**, which were used in solid-phase synthesis of RNA oligonucleotides. Compound **5** was used for internal incorporation of RMP at positions –3 (R-3) and –4 (R-4), and compound **6** was used for synthesis of RNA containing Rem at the 3'-end (R-1).

**Table S1.** Sequences and high-resolution ESI-MS data of synthetic RNA oligonucleotides.

| Name | 5'-sequence-3' | comment | nt | Mass calc. | Mass found |
|---|---|---|---|---|---|
| R-1 | UGAGCCUACGCGR | Prepared with **6** | 13 | 4241.574 Da | 4241.570 Da |
| R-3 | UGAGCCUACGCGRUG | Prepared with **5** | 15 | 4812.680 Da | 4812.702 Da |
| R-4 | UGAGCCUACGCRGUG | Prepared with **5** | 15 | 4812.680 Da | 4812.707 Da |
| A-4 | UGAGCCUACGCAGUG | Unmodified RNA | 15 | 4788.680 Da | 4788.702 Da |

1'-Cyano-5'-*O*-(4,4'-Dimethoxytrityl)-*N*[6]-dimethylformamidine-4-aza-7,9-dideazaadenosine (**2**)

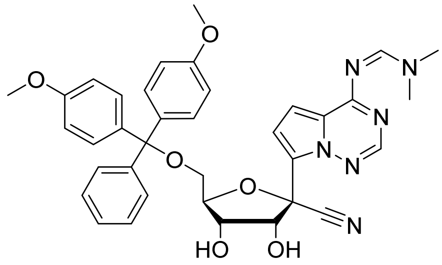

**2**

$C_{36}H_{36}N_6O_6$

648.72 g•mol$^{-1}$

A suspension of 1'-Cyano-4-aza-7,9-dideazaadenosine (**1**, 400 mg, 1.37 mmol, 1.00 eq.) in dry pyridine (4 mL) was treated with *N*,*N*-dimethylformamide dimethyl acetal (550 µL, 4.12 mmol, 3.00 eq.) and stirred at ambient temperature for 18 h. Volatiles were removed *in vacuo* and the residue was redissolved in dry pyridine (4 mL). 4,4'-Dimethoxytrityl chloride (560 mg, 1.65 mmol, 1.20 eq.) was added. The resulting solution was stirred at ambient temperature for 3.5 h. Methanol (16 mL) was added and stirring was continued for 30 min. Volatiles were removed under reduced pressure and the residue was purified by column chromatography (silica gel, $CH_2Cl_2$:MeOH 98:2 + 2% $Et_3N$) to yield the product **2** as a white foam (830 mg, 1.28 mmol, 93%).

**TLC** (silica gel, $CH_2Cl_2$:MeOH 98:2 + 2% $Et_3N$): $R_f$ = 0.40.

**$^1$H NMR** (400 MHz, $CDCl_3$): δ (ppm) = 8.89 (br, 1H, N6-CH), 8.09 (s, 1H, C2-H), 7.27 – 7.23 (m, 2H, trityl-H), 7.21 – 7.13 (m, 7H, trityl-H), 7.04 (d, *J* = 4.6 Hz, 1H, C7-H), 7.00 (d, *J* = 4.6 Hz, 1H, C8-H), 6.77 – 6.70 (m, 4H, trityl-H), 4.78 (d, *J* = 5.3 Hz, 1H, C2'-H), 4.60 (td, *J* = 3.1, 1.7 Hz, 1H, C4'-H), 4.35 (dd, *J* = 5.3, 1.7 Hz, 1H, C3'-H), 3.77 (s, 3H, trityl-$OCH_3$), 3.75 (s, 3H, trityl-$OCH_3$), 3.50 (dd, *J* = 10.6, 3.1 Hz, 1H, C5'-$H^a$), 3.27 (d, J = 0.7 Hz, 3H, $NCH_3^a$), 3.25 (d, J = 0.5 Hz, 3H, $NCH_3^b$), 3.14 (dd, J = 10.6, 3.1 Hz, 1H, C5'-$H^b$).

**$^{13}$C{$^1$H} NMR** (100 MHz, $CDCl_3$): δ (ppm) = 161.07 (1C, C6), 158.56 (2C, trityl-C), 158.08 (1C, N6-CH), 147.91 (1C, C2), 144.57 (1C, trityl-C), 135.87 (1C, trityl-C), 135.51 (1C, trityl-C), 130.13 (2C, trityl-C), 130.09 (2C, trityl-C), 128.14 (2C, trityl-C), 127.91 (2C, trityl-C), 126.90 (1C, trityl-C), 125.69 (1C, C9), 122.53 (1C, C5), 116.78 (1C, CN), 113.23 (2C, trityl-C), 113.20 (2C, trityl-C), 111.46 (1C, C7), 103.87 (1C, C8), 87.20 (1C, C4'), 86.43 (1C, trityl-C), 79.24 (1C, C1'), 77.11 (1C, C2'), 73.49 (1C, C3'), 63.44 (1C, C5'), 55.33 (2C, trityl-$OCH_3$), 55.31 (2C, trityl-$OCH_3$), 41.76 ($NCH_3^a$), 35.54 ($NCH_3^b$).

**HR-ESI-MS**: *m*/*z* calc. ($C_{36}H_{37}N_6O_6$ $[M+H]^+$): 649.27691, found: 649.27836.

1'-Cyano-5'-*O*-(4,4'-Dimethoxytrityl)-*N*[6]-dimethylformamidine-3'-*O*-(*tert*-butyldimethylsilyl)-4-aza-7,9-dideazaadenosine (**3**)

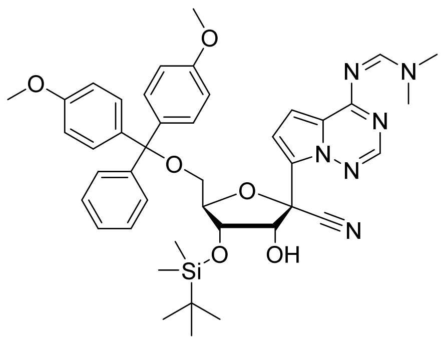

**3**
$C_{42}H_{50}N_6O_6Si$
762.98 g•mol$^{-1}$

A solution of **2** (200 mg, 308 µmol, 1.00 eq.) in dry pyridine (2 mL) was treated with silver nitrate (209 mg, 1.23 mmol, 4.00 eq.) and stirred in the dark at ambient temperature for 30 min. *tert*-Butyldimethylsilyl chloride (55.8 mg, 370 µmol, 1.20 eq.) was added and stirring was continued in the dark for 22 h. Volatiles were removed under reduced pressure. The residue was taken up in ethyl acetate and insoluble residues were removed by filtration through a pad of Celite. The filtrate was evaporated to dryness and the residue was purified by column chromatography (silica gel, *n*-hexane:EtOAc 1:1 + 1% $Et_3N$ to 0:1 + 1% $Et_3N$) to yield the product **3** as a white foam (169 mg, 221 µmol, 72%).

**TLC** (silica gel, *n*-hexane:EtOAc 1:3): $R_f$ = 0.31.

**$^1$H NMR** (400 MHz, $CDCl_3$): δ (ppm) = 8.83 (br, 1H, N6-CH), 8.04 (s, 1H, C2-H), 7.45 – 7.13 (m, 9H, trityl-H), 7.05 (d, *J* = 4.6 Hz, 1H, C7-H), 6.93 (d, *J* = 4.6 Hz, 1H, C8-H), 6.79 – 6.72 (m, 4H, trityl-H), 5.08 (dd, *J* = 9.1, 5.5 Hz, 1H, C2'-H), 4.39 (m, 1H, C4'-H), 4.34 (dd, *J* = 5.5, 2.3 Hz, 1H, C3'-H), 3.78 – 3.75 (m, 7H, (trityl-$OCH_3$)$_2$, C2'-OH), 3.50 (dd, *J* = 10.6, 4.2 Hz, 1H, C5'-H$^a$), 3.25 (d, *J* = 0.6 Hz, 3H, $NCH_3{}^a$), 3.22 (d, *J* = 0.5 Hz, 3H, $NCH_3{}^b$), 3.16 (dd, *J* = 10.6, 3.4 Hz, 1H, C5'-H$^b$), 0.93 (s, 9H, Si-C($CH_3$)$_3$), 0.09 (s, 3H, Si-$CH_3$), 0.00 (s, 3H, Si-$CH_3$).

**$^{13}$C{$^1$H} NMR** (100 MHz, $CDCl_3$): δ (ppm) = 160.73 (1C, C6), 158.46 (2C, trityl-C), 157.55 (1C, N6-CH), 147.38 (1C, C2), 144.51 (1C, trityl-C), 135.84 (1C, trityl-C), 135.59 (1C, trityl-C), 130.02 (4C, trityl-C), 128.15 (2C, trityl-C), 127.79 (2C, trityl-C), 126.80 (1C, trityl-C), 123.60 (1C, C7), 123.45 (1C, C5), 116.53 (1C, CN), 113.08 (4C, trityl-C), 112.69 (1C, C9), 102.52 (1C, C8), 86.38 (1C, trityl-C), 86.21 (1C, C4'), 78.73 (1C, C1'), 74.69 (1C, C2'), 72.77 (1C, C3'), 62.53 (1C, C5'), 55.20 (2C, trityl-$OCH_3$), 41.46 (1C, $NCH_3{}^a$), 35.27 (1C, NCH$^b$), 25.61 (3C, Si-C($\underline{C}H_3$)$_3$), 17.99 (1C, Si-$\underline{C}$($CH_3$)$_3$), -4.63 (1C, Si$CH_3$), -4.99 (1C, Si-$CH_3$).

**HR-ESI-MS**: *m*/*z* calc. ($C_{42}H_{51}N_6O_6Si$ [M+H]$^+$): 763.36339, found: 763.36464.

1'-Cyano-5'-*O*-(4,4'-Dimethoxytrityl)-$N^6$-dimethylformamidine-2'-*O*-(*tert*-butyldimethylsilyl)-4-aza-7,9-dideazaadenosine (**4**)

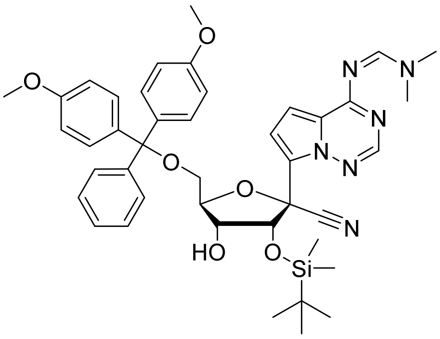

**4**
$C_{42}H_{50}N_6O_6Si$
762.98 g•mol$^{-1}$

Compound **3** (100 mg, 131 µmol, 1.00 eq.) was dissolved in a mixture of methanol (99 mL) and triethylamine (1 mL). After stirring at ambient temperature for 30 min the ratio of isomers **3** and **4** was ca. 1:1 (by TLC). After removal of the solvent under reduced pressure and purification of the residue by column chromatography (*n*-hexane:EtOAc 1:3) 30.0 mg (39.3 µmol, 30%) of pure product **4** were obtained as a white foam. The remaining mixture of isomers was isolated and again treated with trimethylamine in methanol.

**TLC** (silica gel, *n*-hexane:EtOAc 1:3): $R_f$ = 0.41.

**$^1$H NMR** (400 MHz, $CDCl_3$): δ (ppm) = 8.85 – 8.80 (m, 1H, N6-CH), 7.79 (s, 1H, C2-H), 7.51 – 7.43 (m, 2H, trityl-H), 7.39 – 7.32 (m, 4H, trityl-H), 7.28 – 7.15 (m, 3H, trityl-H), 7.11 (d, *J* = 4.6 Hz, 1H, C7-H), 6.91 (d, *J* = 4.6 Hz, 1H, C8-H), 6.82 – 6.73 (m, 4H, trityl-H), 5.50 (d, *J* = 5.7 Hz, 1H, C2'-H), 4.45 (m, 1H, C4'-H), 4.34 (m, 1H, C3'-H), 3.77 (2s, 6H, trityl-$OCH_3$), 3.50 (dd, *J* = 10.4, 4.0 Hz, 1H, C5'-$H^a$), 3.34 (dd, *J* = 10.4, 4.4 Hz, 1H, C5'-$H^b$), 3.26 (d, *J* = 0.7 Hz, 3H, $NCH_3$), 3.22 (d, *J* = 0.4 Hz, 3H, $NCH_3$), 2.77 (d, *J* = 4.3 Hz, 1H, C3'-OH), 0.87 (s, 9H, Si-$C(CH_3)_3$), -0.10 (s, 3H, Si-$CH_3$), -0.15 (s, 3H, Si-$CH_3$).

**$^{13}$C{$^1$H} NMR** (100 MHz, $CDCl_3$): δ (ppm) = 160.83 (1C, C6), 158.55 (1C, trityl-C), 158.53 (1C, trityl-C), 157.71 (1C, N6-CH), 147.20 (1C, C2), 145.03 (1C, trityl-C), 136.22 (1C, trityl-C), 136.11 (1C, trityl-C), 130.28 (4C, trityl-C), 128.41 (2C, trityl-C), 127.87 (2C, trityl-C), 126.85 (1C, trityl-C), 123.86 (1C, C5), 122.29 (1C, C7), 117.10 (1C, CN), 114.81 (1C, C9), 113.18 (4C, trityl-C), 102.52 (1C, C8), 86.25 (1C, trityl-C), 85.43 (1C, C4'), 80.06 (1C, C1'), 73.61 (1C, C2'), 71.68 (1C, C3'), 63.02 (1C, C5'), 55.34 (2C, trityl-$OCH_3$), 41.62 (1C, $NCH_3$), 35.42 (1C, $NCH_3$), 25.75 (3C, Si-$C(\underline{C}H_3)_3$), 18.11 (1C, Si-$\underline{C}(CH_3)_3$), -4.91 (1C, Si-$CH_3$), -5.12 (1C, Si-$CH_3$).

**HR-ESI-MS**: *m*/*z* calc. ($C_{42}H_{51}N_6O_6Si$ $[M+H]^+$): 763.36339, found: 763.36413.

1'-Cyano-5'-*O*-(4,4'-Dimethoxytrityl)-*N*[6]-dimethylformamidine-2'-*O*-(*tert*-butyldimethylsilyl)-4-aza-7,9-dideazaadenosine 3'-β-cyanoethyl diisopropyl phosphoramidite (**5**)

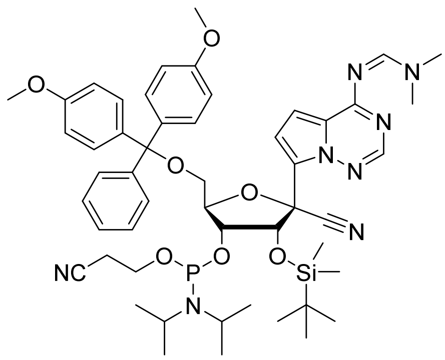

**5**

$C_{51}H_{67}N_8O_7PSi$

963.20 g•mol$^{-1}$

A solution of **4** (110.0 mg, 144 µmol, 1.00 eq.) in dry dichloromethane (1.1 mL) was treated with *N*,*N*-dimethylethylamine (156 µL, 1.44 mmol, 10.0 eq.) and 2-cyanoethyl *N*,*N*-diisopropyl-chlorophosphoramidite (40.9 mg, 173 µmol, 1.20 eq.) and stirred at ambient temperature for 5 h. Volatiles were removed under reduced pressure and the residue was purified by column chromatography (silica gel, *n*-hexane:EtOAc 1:3) to yield the desired product as a white foam (104 mg, 108 µmol, 75%, isomer ratio at phosphorus 10:8).

**TLC** (silica gel, *n*-hexane:EtOAc 1:3): $R_f$ = 0.45.

**$^1$H NMR** (400 MHz, $CDCl_3$): δ (ppm) = 8.82, 8.81 (2 s, N6-CH), 7.68 (s, C2-H), 7.58 (s, C2-H), 7.54 – 7.45 (m, trityl-H), 7.43 – 7.33 (m, trityl-H), 7.27 – 7.18 (m, trityl-H), 7.16, 7.15 (2 d, *J* = 4.6 Hz, C7-H), 6.92, 6.91 (2 d, *J* = 4.6 Hz, C8-H), 6.83 – 6.72 (m, trityl-H), 5.63, 5.61 (2 d, *J* = 5.3 Hz, C2'-H), 4.62 (app q, *J* = 3.3 Hz, C4'-H), 4.58 – 4.51 (m, C4'-H), 4.45 – 4.34 (m, C3'-H), 4.20 – 4.09 (m, $POCH_2$), 3.98 – 3.88 (m, $POCH_2$), 3.78 – 3.75 (m, trityl-$OCH_3$), 3.66 – 3.52 (m, C5'-H, $N(C\underline{H}(CH_3)_2)_2$), 3.34 (dd, *J* = 10.3, 4.5 Hz, C5'-H), 3.28 – 3.23 (m, C5'-H, $N(CH_3)_2$), 3.21 (d, *J* = 0.9 Hz, $N(CH_3)_2$), 2.73 – 2.54 (m, $CH_2CN$), 2.24 – 2.18 (m, $CH_2CN$), 1.18 (d, *J* = 6.8 Hz, $N(CH(C\underline{H}_3)_2)_2$), 1.15 (d, *J* = 6.8 Hz, $N(CH(C\underline{H}_3)_2)_2$), 1.00 (d, *J* = 6.8 Hz, $N(CH(C\underline{H}_3)_2)_2$), 0.80 (s, 4H), 0.77 (s, $SiCH_3$), -0.09 (s, $SiCH_3$), -0.18 (s, $SiCH_3$), -0.33 (s, $SiCH_3$), -0.40 (s, $SiCH_3$).

**$^{13}$C{$^1$H} NMR** (100 MHz, $CDCl_3$): δ (ppm) = 160.79 (C6), 158.55 (trityl-C), 157.67 (N6-CH), 147.07 (C2), 146.98 (C2), 145.05 (trityl-C), 144.87 (trityl-C), 136.29 (trityl-C), 136.17 (trityl-C), 136.09 (trityl-C), 136.02 (trityl-C), 130.41 (trityl-C), 130.36 (trityl-C), 128.57 (trityl-C), 128.52 (trityl-C), 127.85 (trityl-C), 126.88 (trityl-C), 123.98 (C5), 123.81 (C5), 122.73 (C9), 122.22 (C9), 118.34 ($CH_2\underline{C}N$), 117.95 (C1'-$\underline{C}N$), 117.53 ($CH_2\underline{C}N$), 117.32 (C1'-$\underline{C}N$), 116.00 (C7), 115.46 (C7), 113.15 (trityl-C), 113.13 (trityl-C), 102.51 (C8), 86.43 (trityl-C), 86.24 (trityl-C), 85.79 (C4'), 79.90 (C1'), 79.69 (C1'), 73.45 (C2'), 72.46 (C3'), 72.35 (C3'), 63.32 (C5'), 63.04 (C5'), 59.26 ($POCH_2$), 59.14 ($POCH_2$), 57.78 ($POCH_2$), 57.59 ($POCH_2$), 55.35 (trityl-$OCH_3$), 43.55 ($N\underline{C}H(CH_3)_2$), 43.43 ($N\underline{C}H(CH_3)_2$), 43.04 ($N\underline{C}H(CH_3)_2$), 42.91 ($N\underline{C}H(CH_3)_2$), 41.59 ($NCH_3$), 35.41 ($NCH_3$), 25.87 ($SiCH(\underline{C}H_3)_3$) 25.85 ($SiCH(\underline{C}H_3)_3$), 25.81 ($SiCH(\underline{C}H_3)_3$), 24.93 ($NCH(\underline{C}H_3)_2$), 24.85 ($NCH(\underline{C}H_3)_2$), 24.75 ($NCH(\underline{C}H_3)_2$), 24.69 ($NCH(\underline{C}H_3)_2$), 24.63 ($NCH(\underline{C}H_3)_2$), 20.88 ($\underline{C}H_2$-CN), 20.83 ($\underline{C}H_2$-CN), 18.11 ($Si\underline{C}H(CH_3)_3$), 18.04 ($Si\underline{C}H(CH_3)_3$), -4.64 ($SiCH_3$), -4.68 ($SiCH_3$), -5.37 ($SiCH_3$), -5.45 ($SiCH_3$).

**$^{31}$P{$^1$H} NMR** (162 MHz, $CDCl_3$): δ (ppm) = 150.48, 147.94.

**HR-ESI-MS**: *m*/*z* calc. ($C_{51}H_{68}N_8O_7PSi$ $[M+H]^+$): 963.47124, found: 963.46940.

1'-Cyano-5'-*O*-(4,4'-Dimethoxytrityl)-$N^6$-dimethylformamidine-3'-*O*-(*tert*-butyldimethylsilyl)-4-aza-7,9-dideazaadenosine 2'-β-cyanoethyl diisopropyl phosphoramidite (**6**)

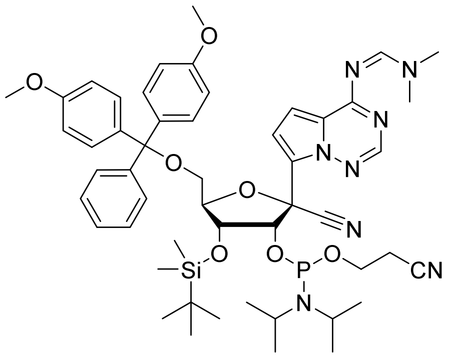

**6**
$C_{51}H_{67}N_8O_7PSi$
963.20 g•mol$^{-1}$

A solution of **3** (50.0 mg, 65.5 µmol, 1.00 eq.) in dry dichloromethane (0.5 mL) was treated with *N*,*N*-dimethylethylamine (71.0 µL, 655 µmol, 10.0 eq.) and 2-cyanoethyl *N*,*N*-diisopropyl-chlorophosphoramidite (18.6 mg, 78.6 µmol, 1.20 eq.) and stirred at ambient temperature for 5 h. Volatiles were removed under reduced pressure and the residue was purified by column chromatography (silica gel, *n*-hexane:EtOAc 1:2) to yield the product as a white foam (54.0 mg, 56.1 µmol, 85%, isomer ratio at phosphorus 10:3).

**TLC** (silica gel, *n*-hexane:EtOAc 1:2): $R_f$ = 0.20.

**$^1$H NMR** (400 MHz, $CDCl_3$): δ (ppm) = 8.82 (s, N6-CH), 8.79 (t, *J* = 0.7 Hz, N6-CH), 7.92 (s, C2-H), 7.72 (s, C2-H), 7.46 – 7.16 (m, C7-H, trityl-H), 7.13 (dd, *J* = 4.6, 0.8 Hz, C7-H), 6.91 (d, *J* = 4.6 Hz, C8-H), 6.86 (d, *J* = 4.6 Hz, C8-H), 6.82 – 6.74 (m, trityl-H), 5.37 (dd, *J* = 12.6, 4.7 Hz, 1H, C2'-H), 5.28 (dd, *J* = 10.8, 4.1 Hz, 1H, C2'-H), 4.48 – 4.37 (m, 3'-H, 4'-H), 3.90 – 3.82 (m, $POCH_2$), 3.82 – 3.72 (m, (trityl-$OCH_3$)$_2$, $POCH_2$), 3.69 – 3.47 (m, $POCH_2$, N(C$\underline{H}$($CH_3$)$_2$)$_2$, C5'-H), 3.30 (dd, *J* = 10.2, 4.1 Hz, C5'-H), 3.26 – 3.19 (m, N($CH_3$)$_2$, C5'-H), 2.78 – 2.31 (m, $CH_2CN$), 1.11 (d, *J* = 6.8 Hz, N(CH(C$\underline{H}_3$)$_2$)$_2$), 1.05 (d, *J* = 6.7 Hz, N(CH(C$\underline{H}_3$)$_2$)$_2$), 1.02 – 0.94 (m, N(CH(C$\underline{H}_3$)$_2$)$_2$, Si-C($CH_3$)$_3$), 0.91 (s, Si-C($CH_3$)$_3$), 0.14 (s, Si-$CH_3$), 0.12 (s, Si-$CH_3$), 0.09 (s, Si-$CH_3$), 0.00 (s, Si-$CH_3$).

**$^{13}$C{$^1$H} NMR** (100 MHz, $CDCl_3$): δ (ppm) = 160.78 (C6), 158.58 (trityl-C), 158.56 (trityl-C), 157.63 (N6-CH), 147.11 (C2), 144.82 (trityl-C), 136.07 (trityl-C), 135.92 (trityl-C), 130.28 (trityl-C), 130.24 (trityl-C), 130.16 (trityl-C), 128.46 (trityl-C), 128.40 (trityl-C), 127.93 (trityl-C), 127.89 (trityl-C), 126.92 (trityl-C), 123.65 (C5), 123.42 (C9), 117.75 ($CH_2\underline{C}N$), 117.07 (C1'-$\underline{C}$N), 114.03 (C7), 113.99 (C7), 113.21 (trityl-C), 113.19 (trityl-C), 102.44 (C8), 86.82 (trityl-C), 86.60 (trityl-C), 86.45 (C4'), 86.33 (C4'), 77.87 (C1'), 77.82 (C1'), 75.18 (C2'), 75.04 (C2'), 72.86 (C3'), 63.23 (C5'), 58.64 ($POCH_2$), 58.47 ($POCH_2$), 55.37 (trityl-$OCH_3$), 43.41 (N($\underline{C}$H($CH_3$)$_2$)$_2$), 43.29 (N($\underline{C}$H($CH_3$)$_2$)$_2$), 41.59 ($NCH_3$), 35.39 ($NCH_3$), 25.85 (Si-C($\underline{C}H_3$)$_3$), 24.83 (N(CH($\underline{C}H_3$)$_2$)$_2$), 24.76 (N(CH($\underline{C}H_3$)$_2$)$_2$), 24.69 (N(CH($\underline{C}H_3$)$_2$)$_2$), 24.61 (N(CH($\underline{C}H_3$)$_2$)$_2$), 24.39 (N(CH($\underline{C}H_3$)$_2$)$_2$), 24.32 (N(CH($\underline{C}H_3$)$_2$)$_2$), 20.46 ($\underline{C}H_2CN$), 20.40 ($\underline{C}H_2CN$), 18.11 (Si-$\underline{C}$($CH_3$)$_3$), -4.33 ($SiCH_3$), -4.36 ($SiCH_3$), -4.40 ($SiCH_3$), -4.42 ($SiCH_3$).

**$^{31}$P{$^1$H} NMR** (162 MHz, $CDCl_3$): δ (ppm) = 149.92, 149.25.

**HR-ESI-MS**: *m*/*z* calc. ($C_{51}H_{68}N_8O_7PSi$ [M+H]$^+$): 963.47124, found: 963.47136.

### NMR Spectra

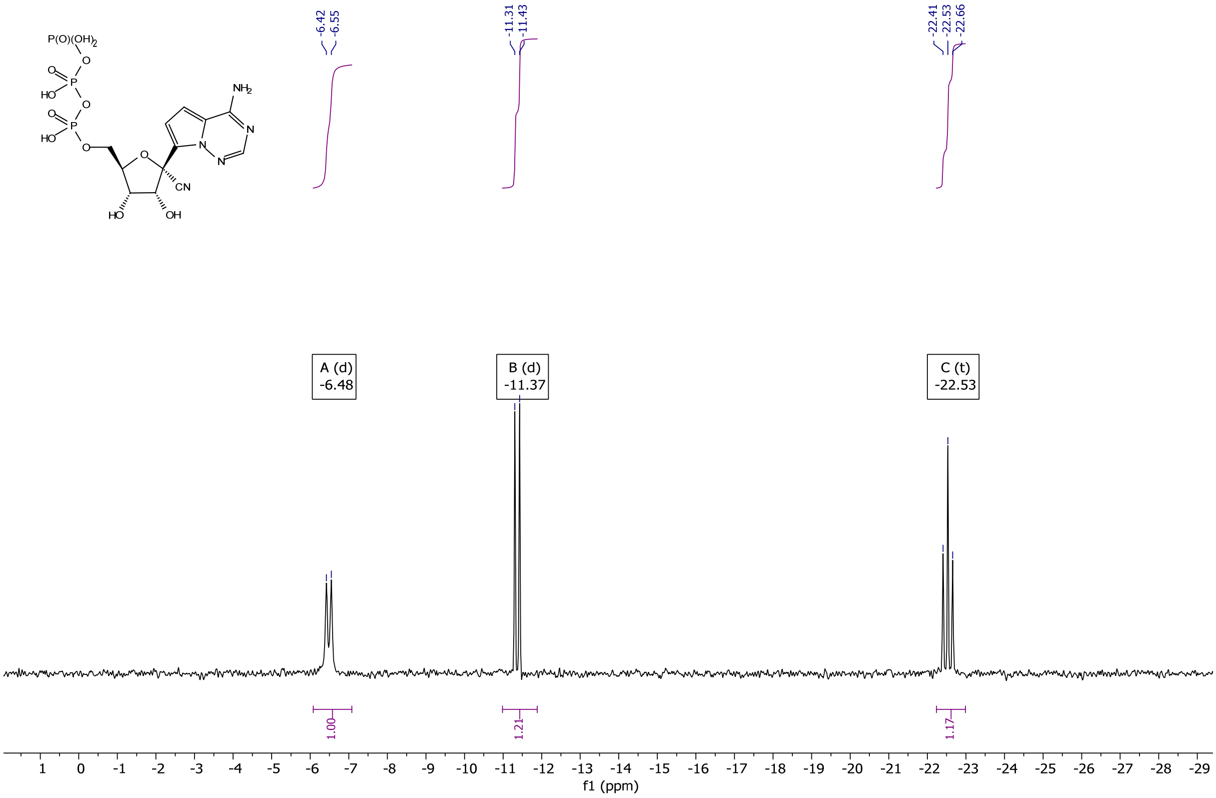

$^{31}P$ NMR (162 MHz, $D_2O$) of **RTP***5$(C_2H_5)_3N$

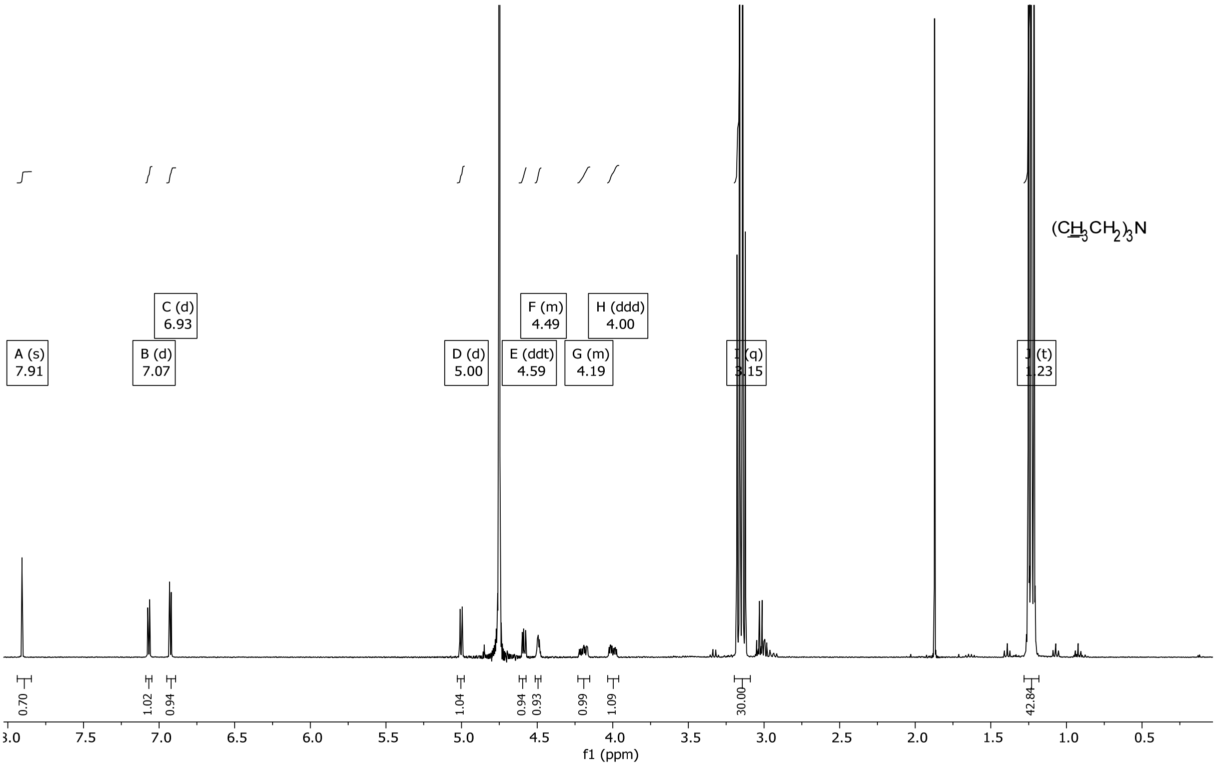

$^{1}H$ NMR (400 MHz, $D_2O$) of **RTP*** 5$(C_2H_5)_3N$

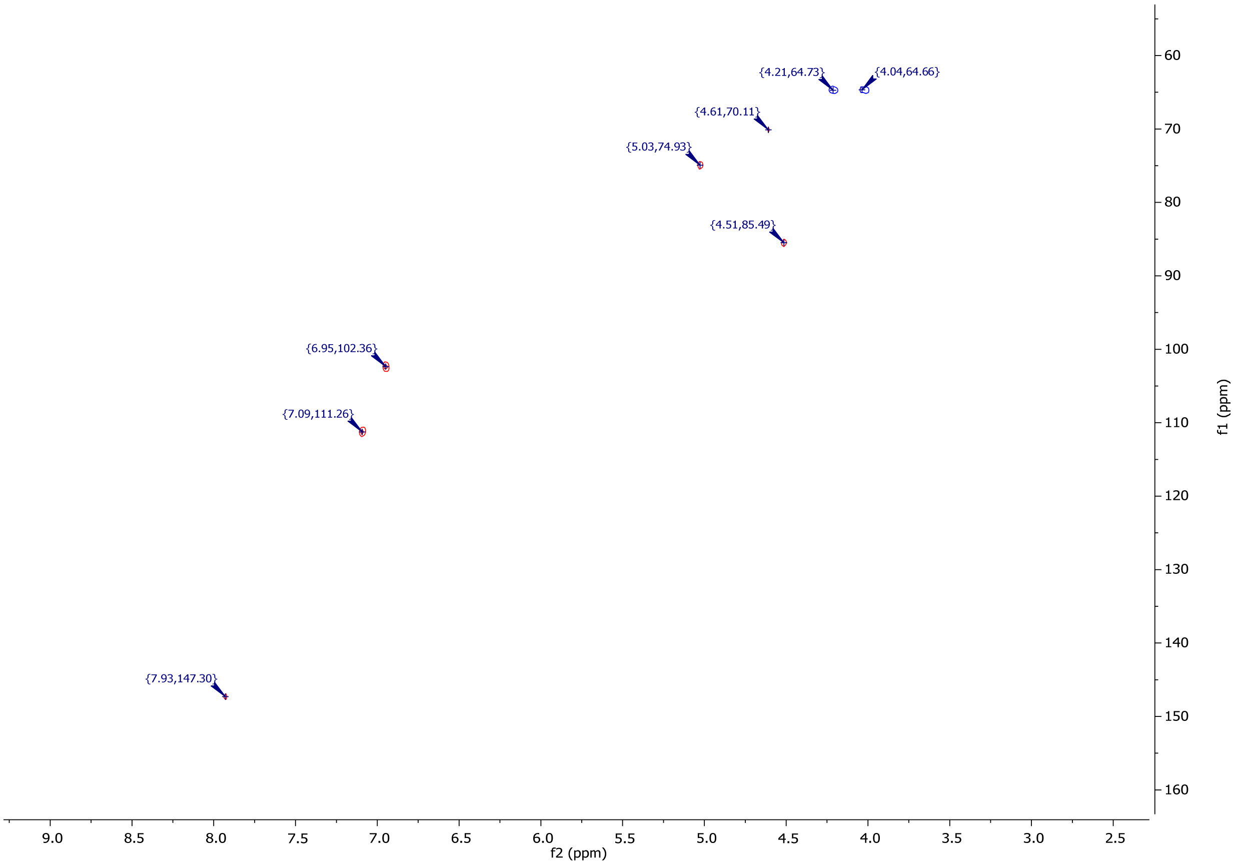

$^{1}H$ /$^{13}C$-HSQC NMR (400 MHz, $D_2O$) of **RTP***5$(C_2H_5)_3N$

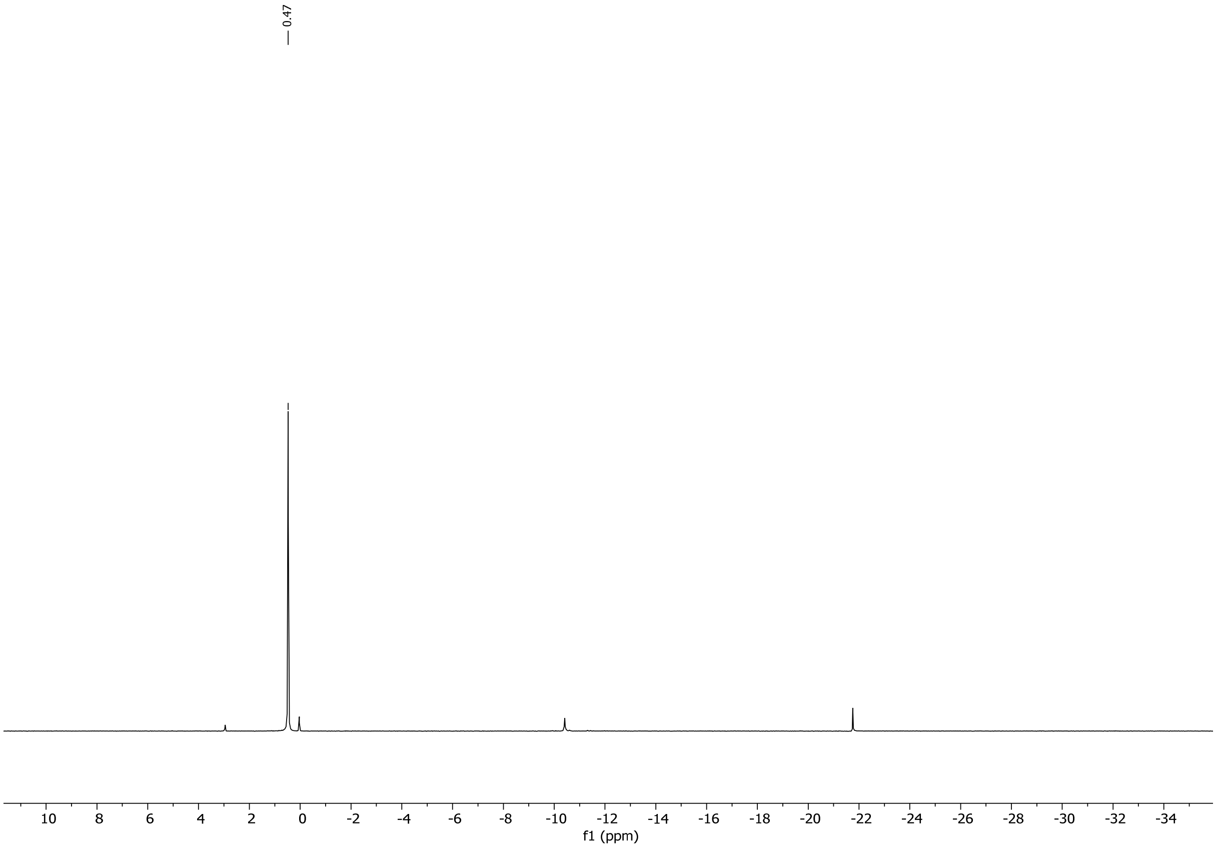

$^{31}P$ NMR (162 MHz, $D_2O$) of **RMP**.

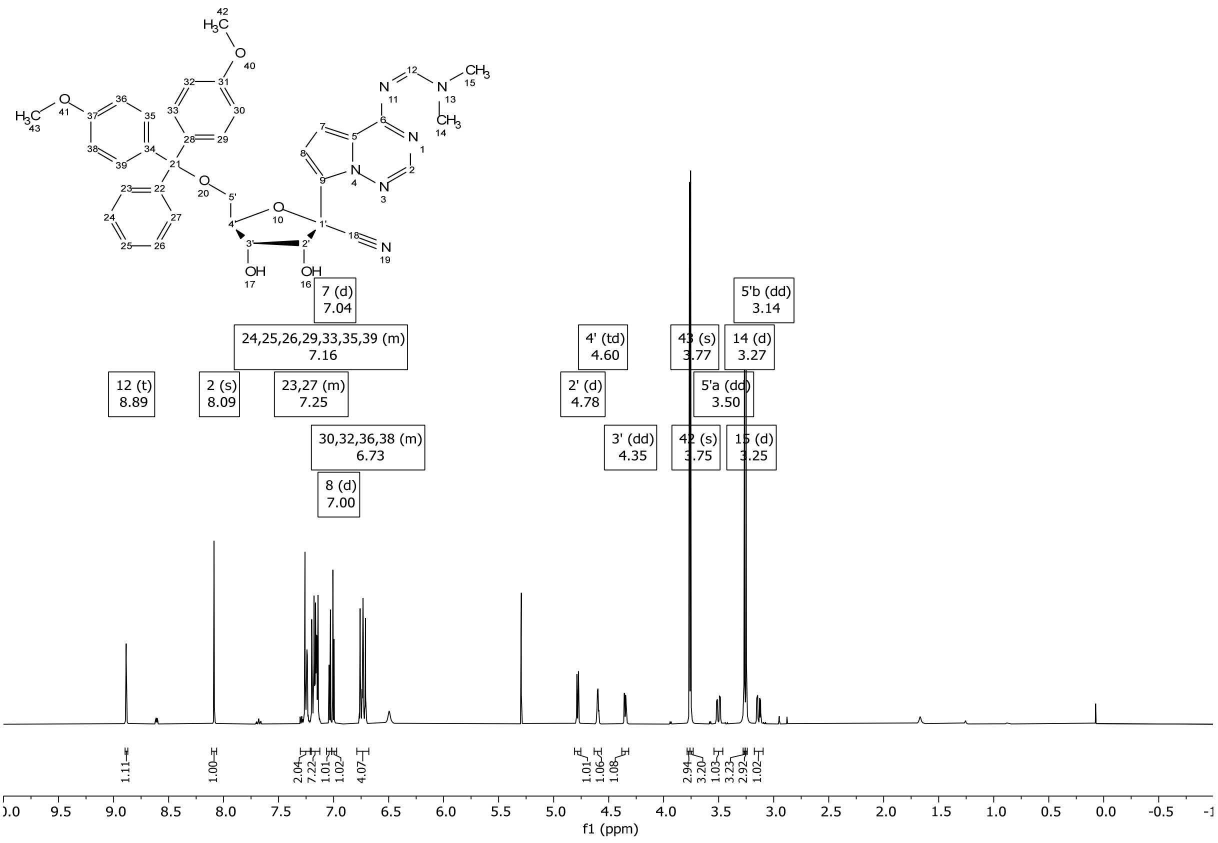

$^{1}H$ NMR (400 MHz, $CDCl_3$) of compound **2**.

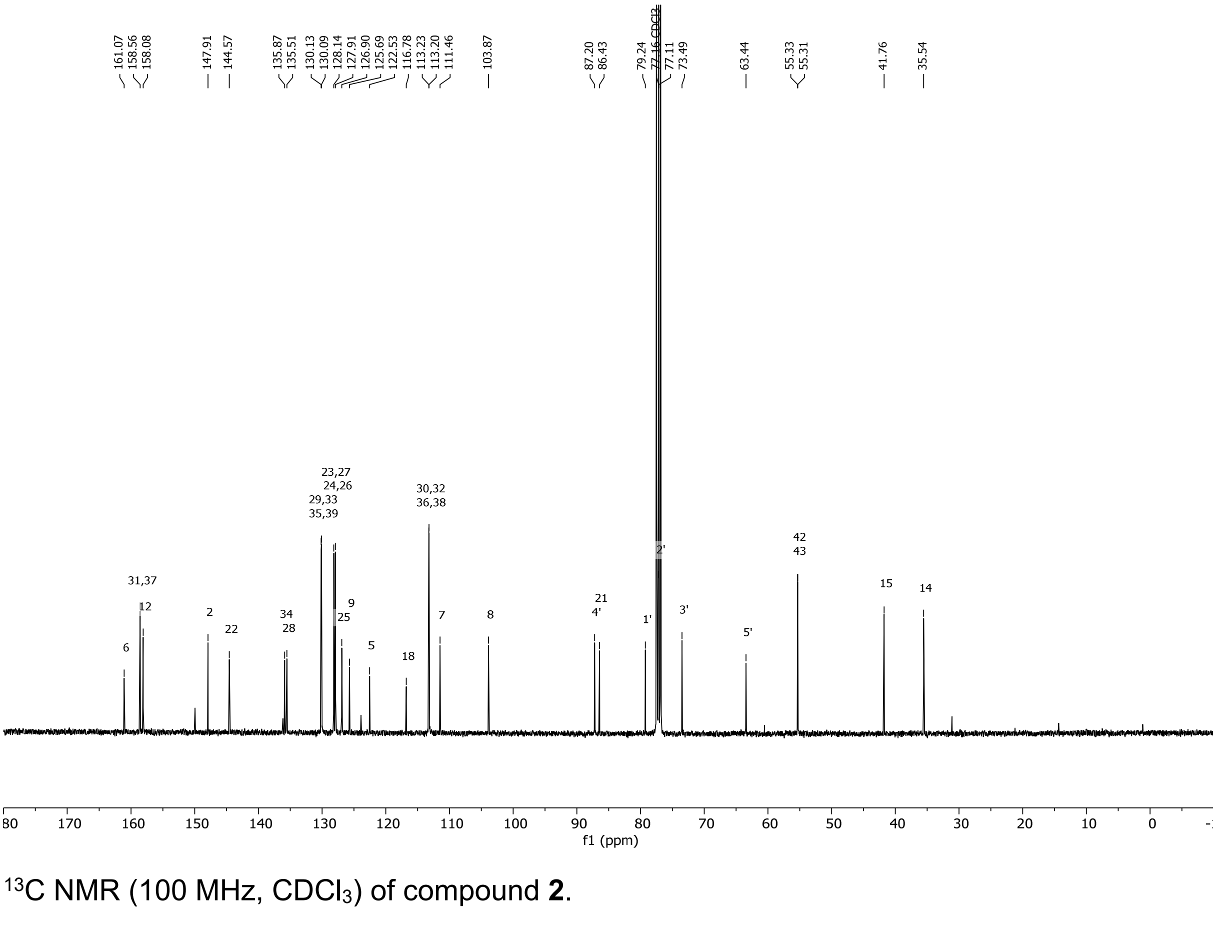

$^{13}C$ NMR (100 MHz, $CDCl_3$) of compound **2**.

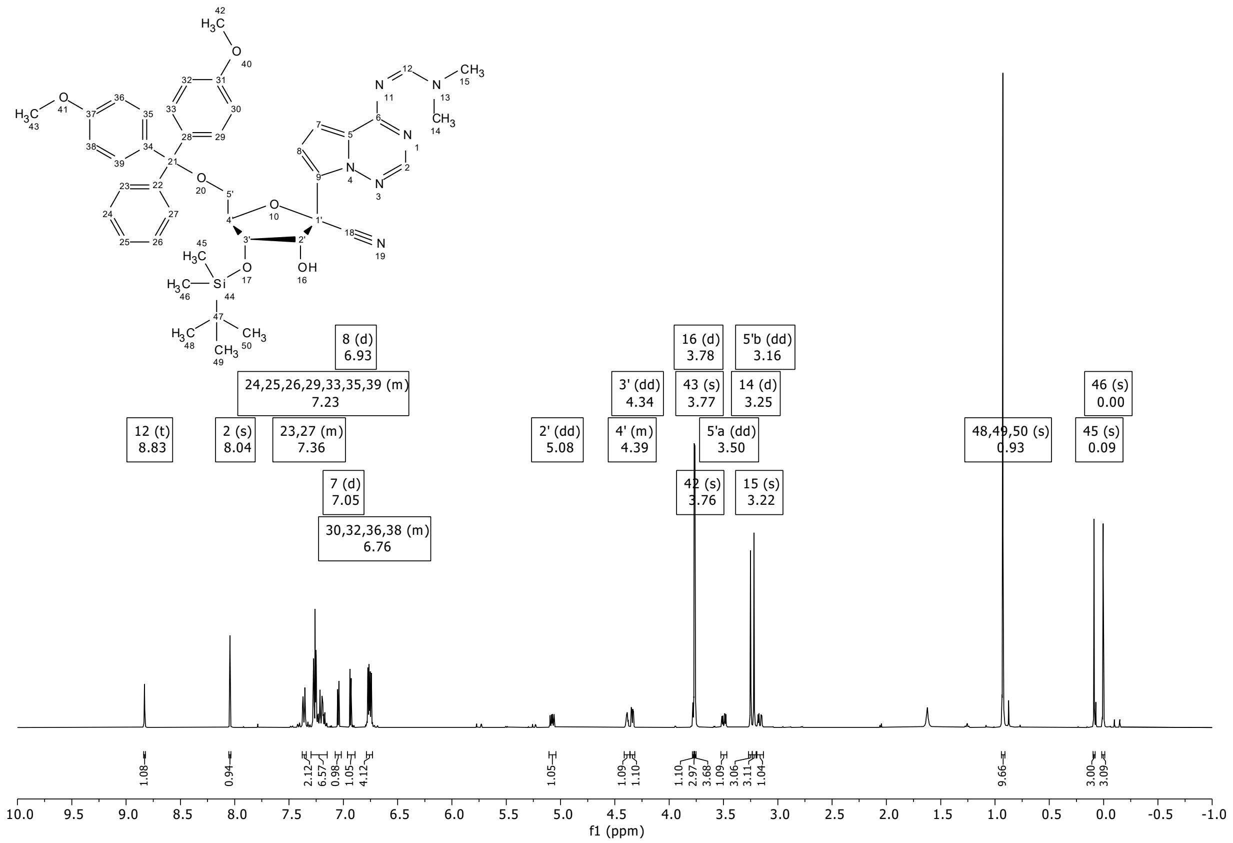

$^{1}H$ NMR (400 MHz, $CDCl_3$) of compound **3**.

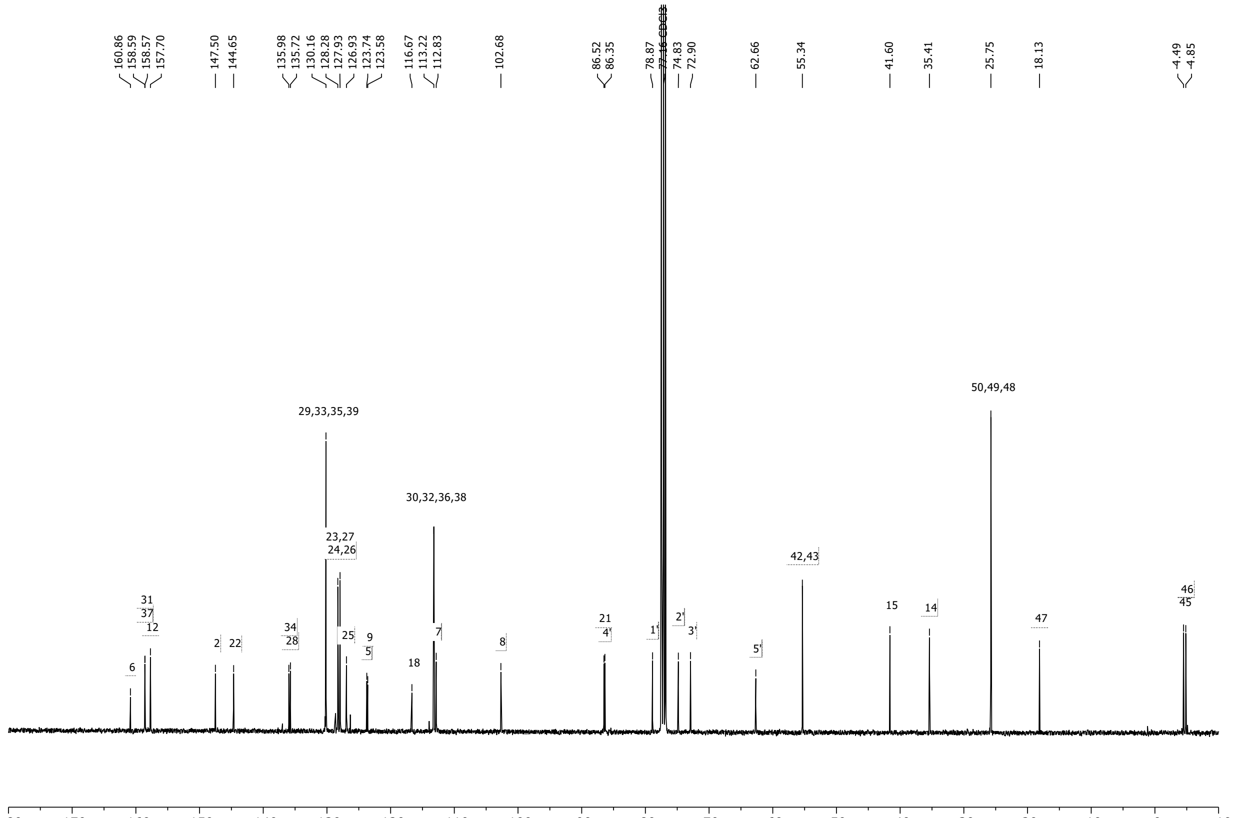

$^{13}C$ NMR (100 MHz, $CDCl_3$) of compound **3**.

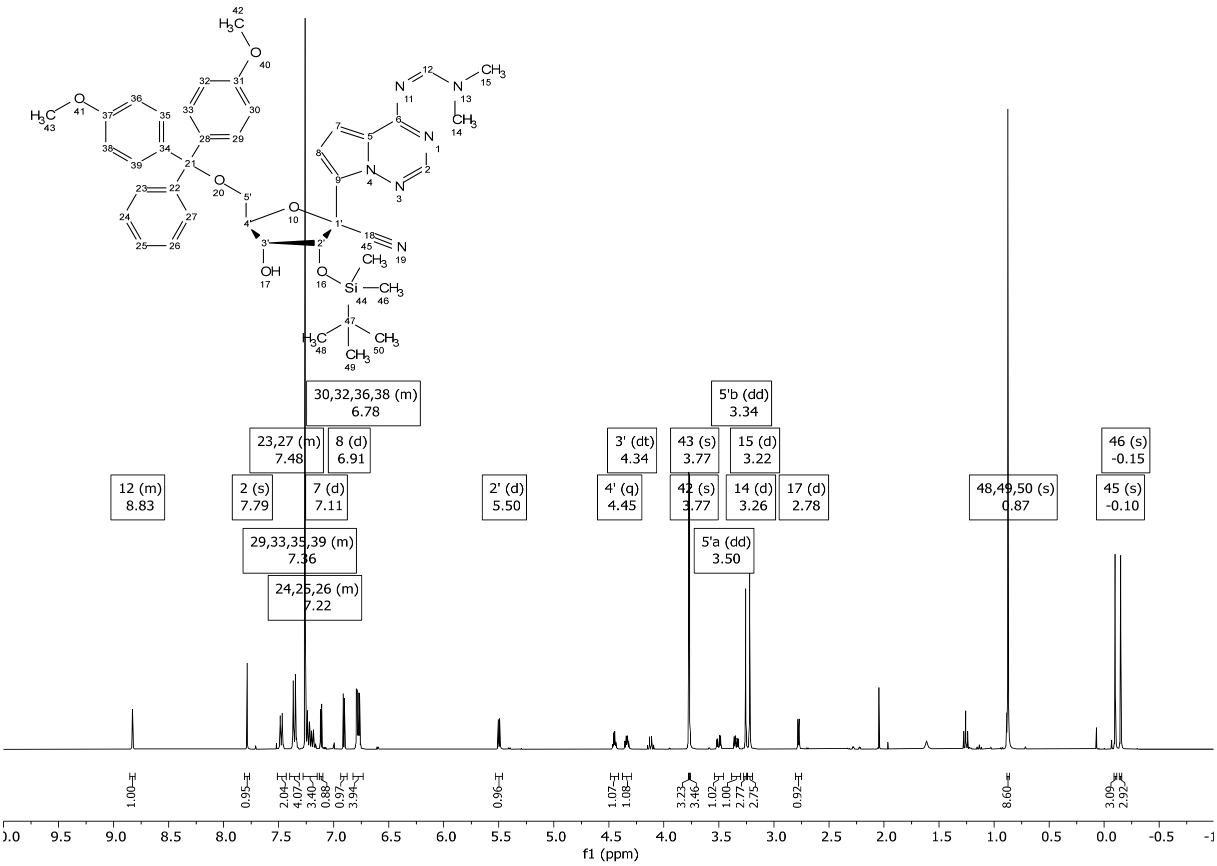

¹H NMR (400 MHz, $CDCl_3$) of compound **4**.

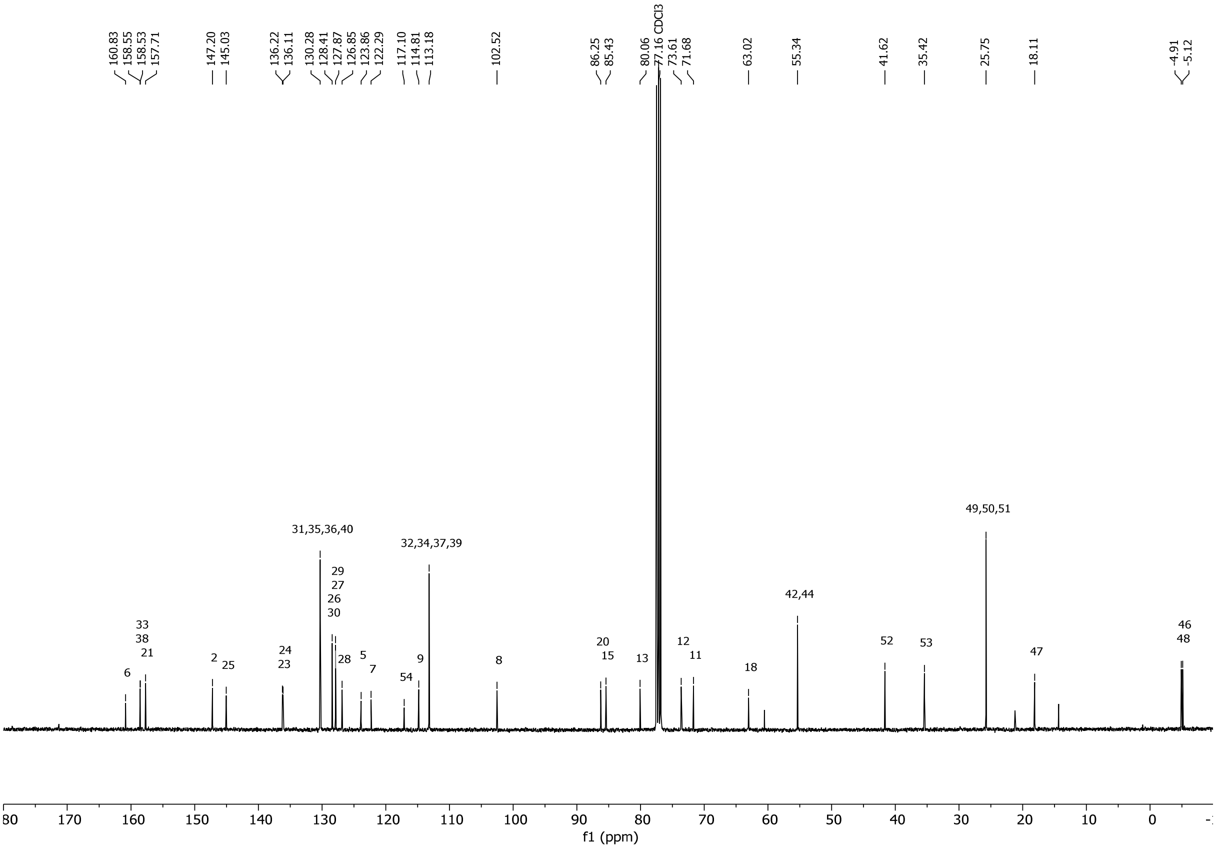

¹³C NMR (100 MHz, $CDCl_3$)of compound **4**.

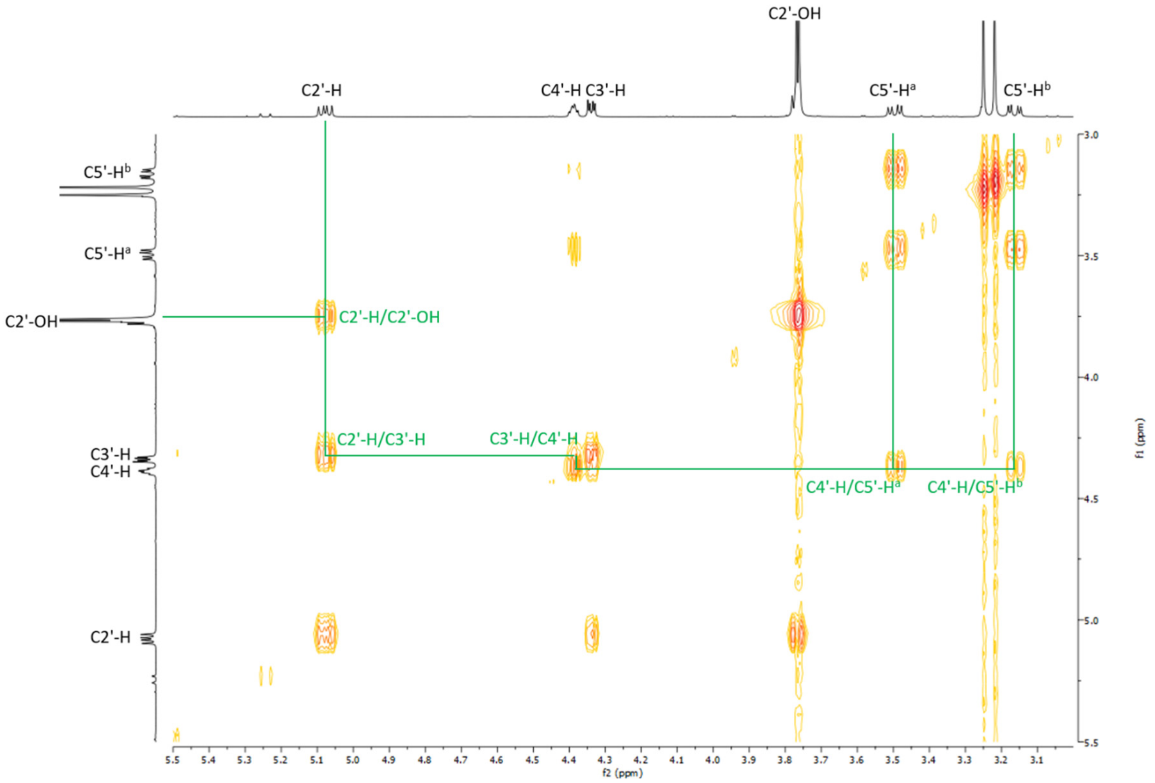

Excerpt of the [1]H/[1]H-COSY NMR spectrum of compound **3**. The relevant cross-peaks displaying the connectivity of the ribose protons are highlighted. The presence of the C2'-H/C2'-OH cross-peak confirms that the TBDMS protecting group of **3** is attached to the C3'-OH.

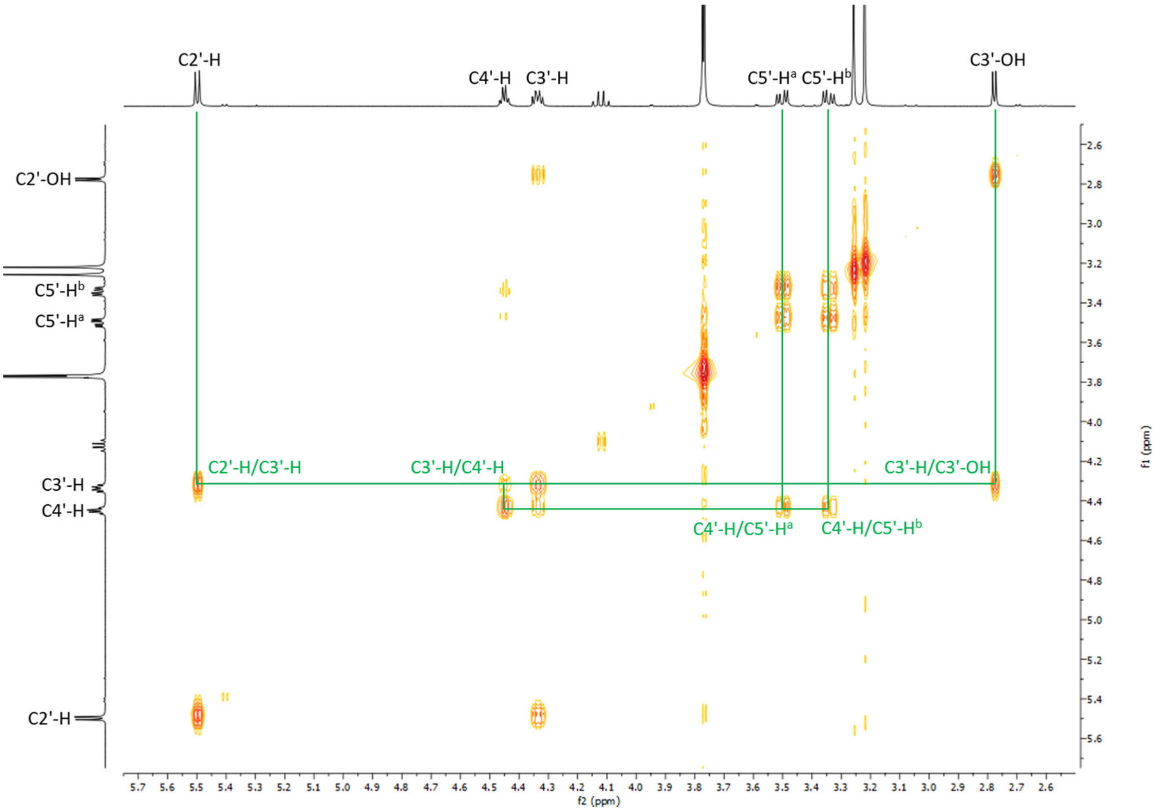

Excerpt of the [1]H/[1]H-COSY NMR spectrum of compound **4**. The relevant cross-peaks displaying the connectivity of the ribose protons are highlighted. The presence of the C3'-H/C3'-OH cross-peak confirms that the TBDMS protecting group of **4** is attached to the C2'-OH.

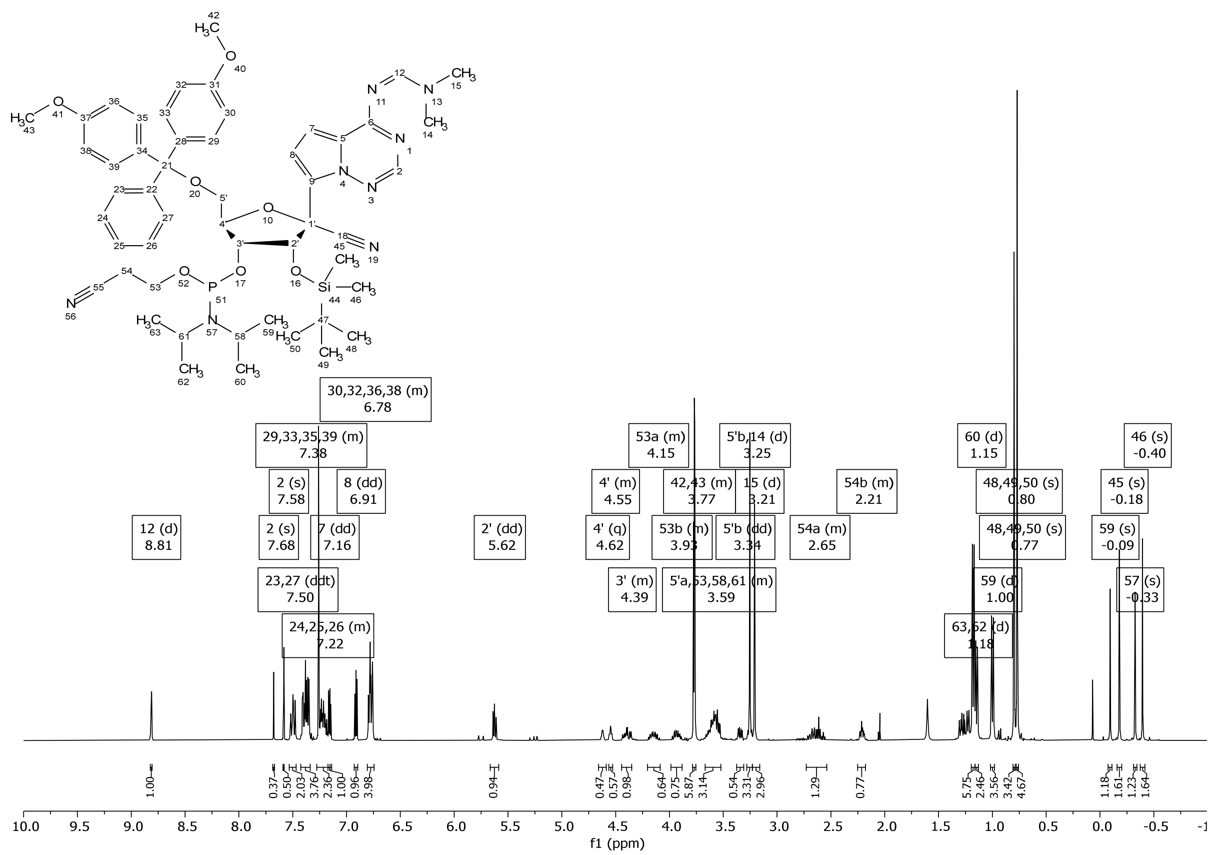

[1]H NMR (400 MHz, $CDCl_3$) of compound **5**.

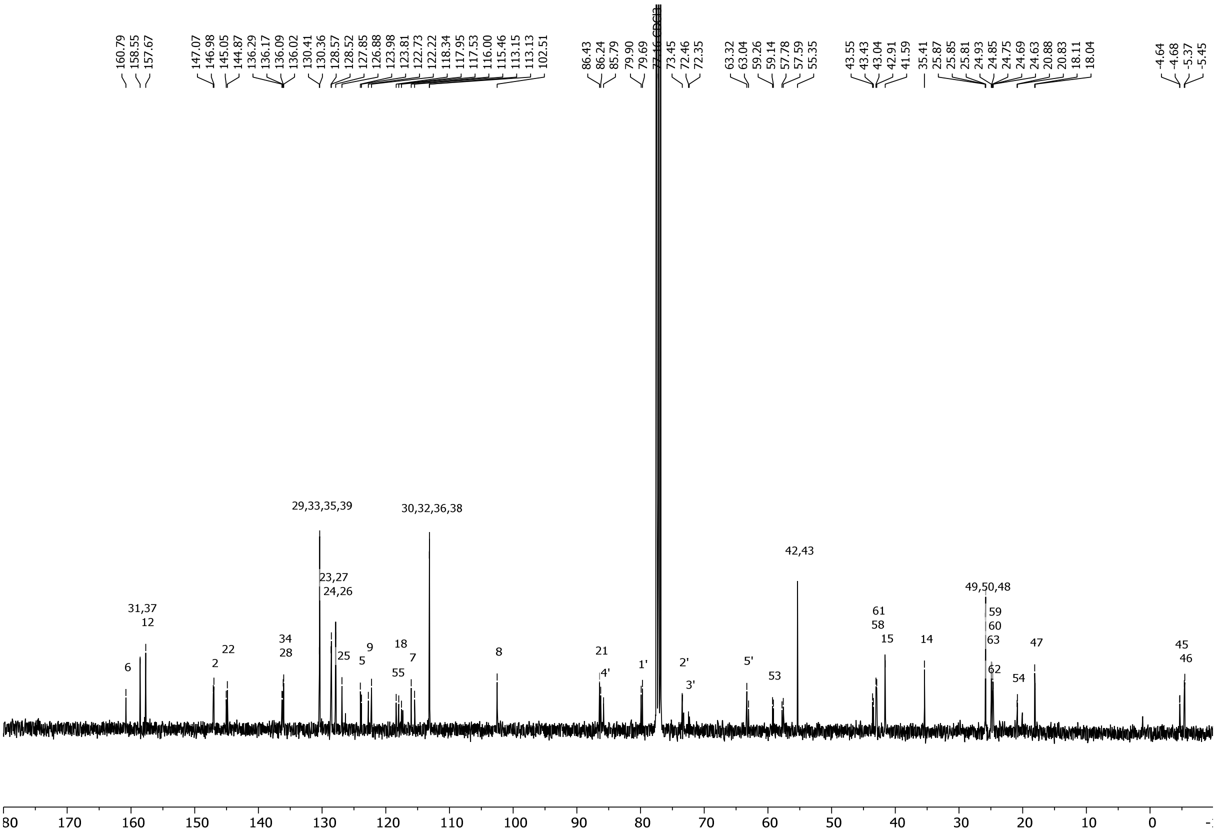

[13]C NMR (100 MHz, $CDCl_3$) of compound **5**.

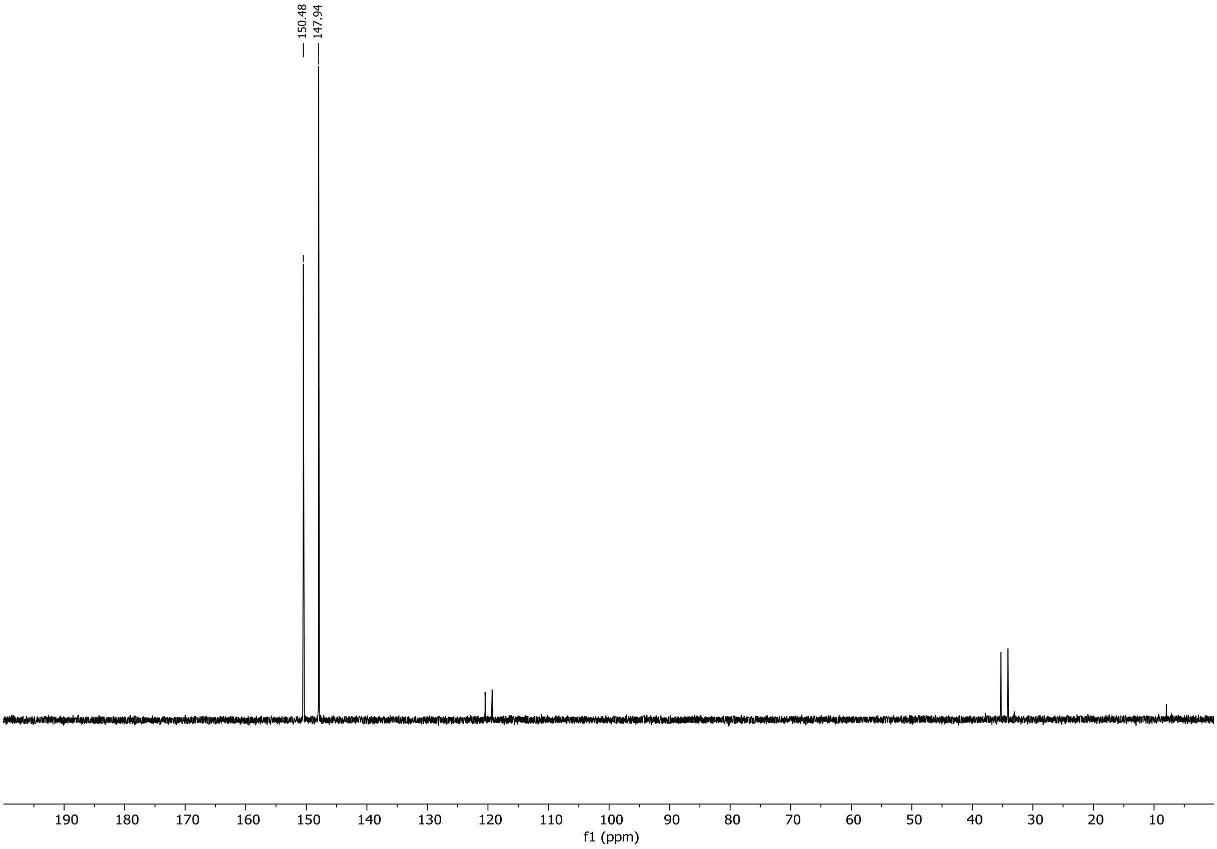

$^{31}$P NMR (162 MHz, $CDCl_3$) of compound **5**.

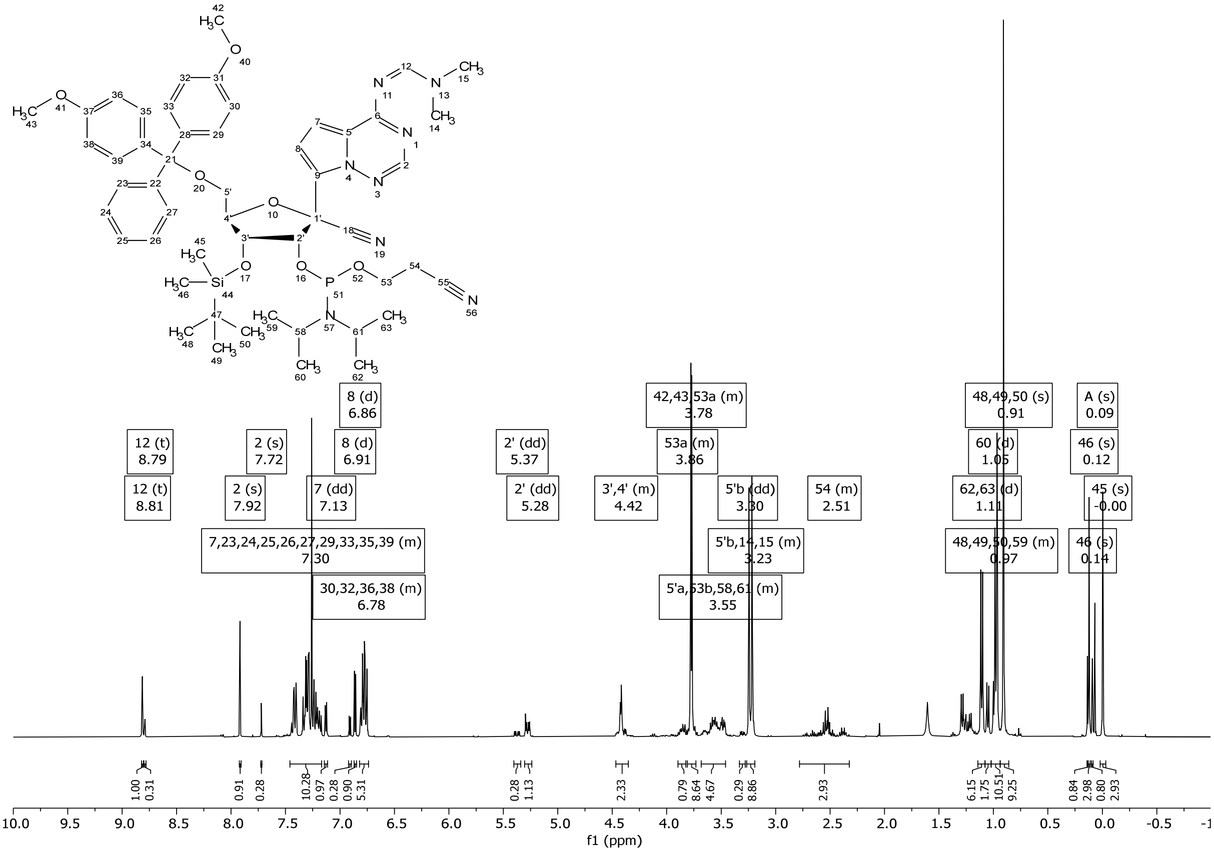

$^{1}$H NMR (400 MHz, $CDCl_3$) of compound **6**.

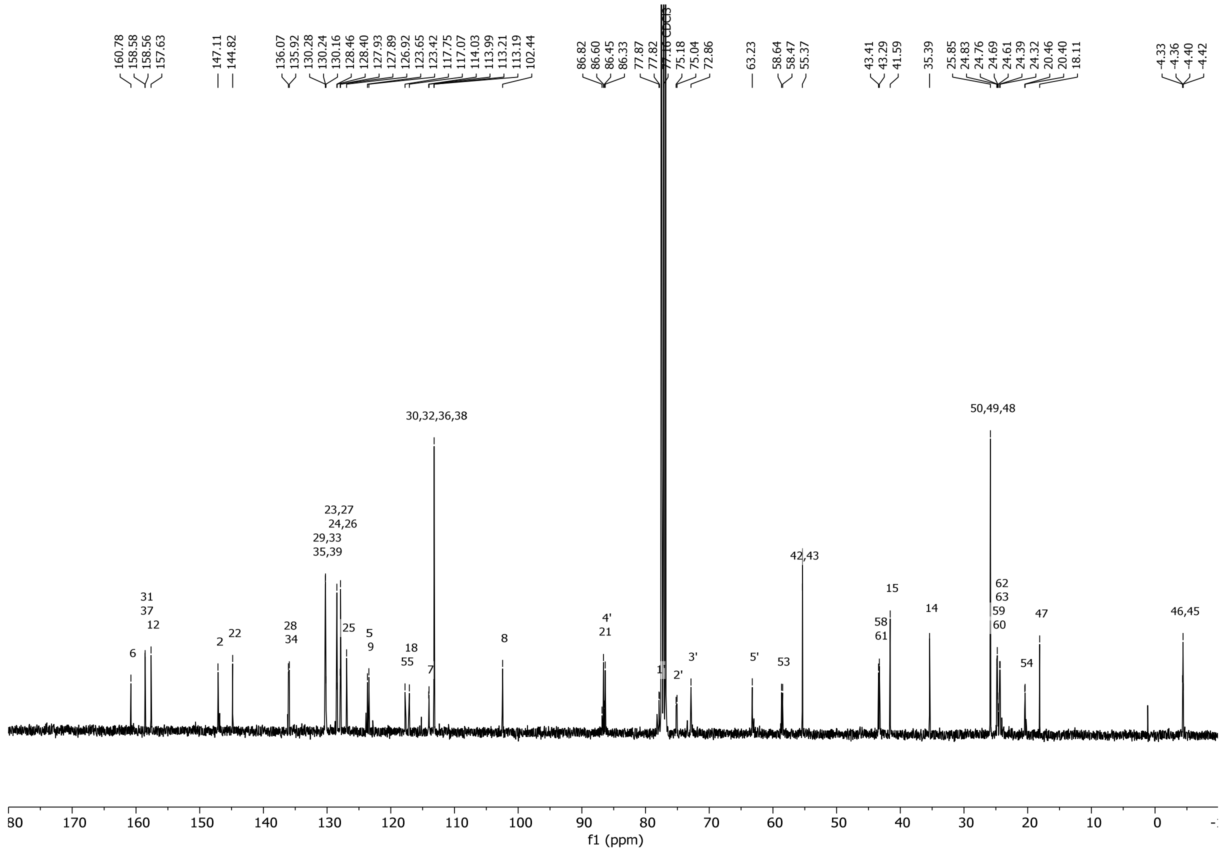

$^{13}C$ NMR (100 MHz, $CDCl_3$) of compound **6**.

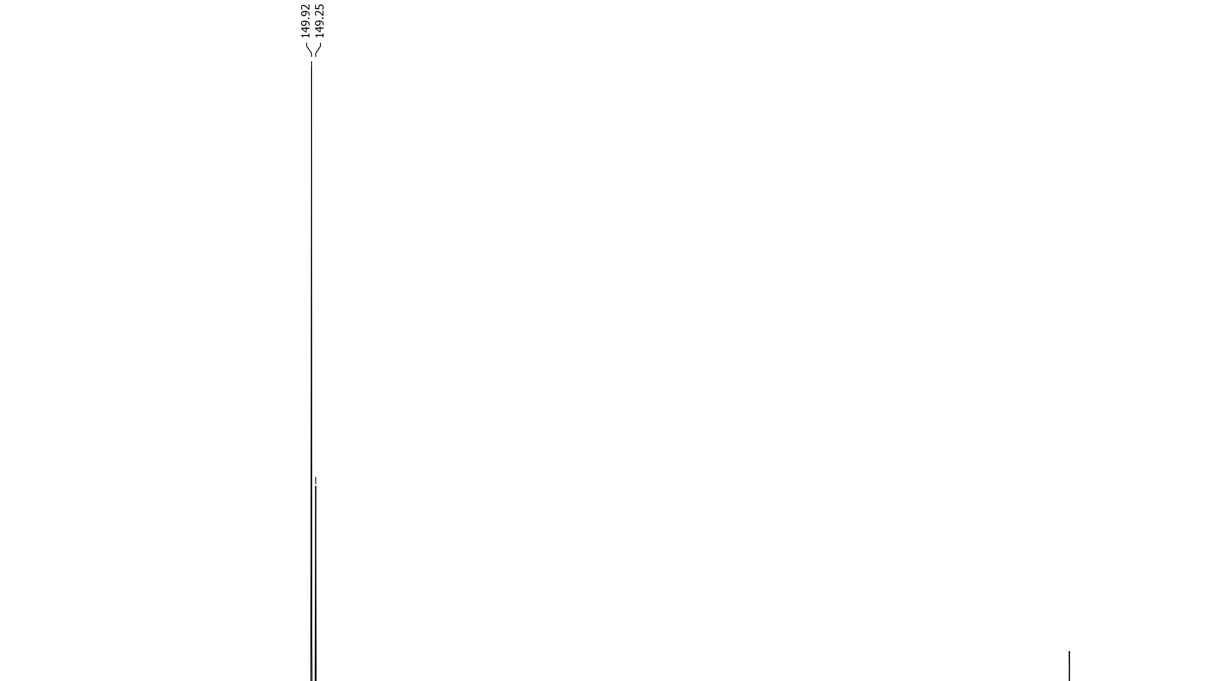

$^{31}P$ NMR (162 MHz, $CDCl_3$) of compound **6**.
